# Single-cell RNA-seq identifies a reversible epithelial-mesenchymal transition in abnormally specified epithelia of p63 EEC syndrome

**DOI:** 10.1101/437632

**Authors:** Eduardo Soares, Quan Xu, Qingqing Li, Jieqiong Qu, Yuxuan Zheng, Henriette H. M. Raeven, Karina Brandao, Isabelle Petit, Willem M.R. van den Akker, Daniel Aberdam, Fuchou Tang, Huiqing Zhou

## Abstract

Mutations in transcription factor p63 are associated with developmental disorders that manifest defects in stratified epithelia including the epidermis. The underlying cellular and molecular mechanism is however not yet understood. We established an epidermal commitment model using human induced pluripotent stem cells (iPSCs) and characterized differentiation defects of iPSCs derived from ectrodactyly, ectodermal dysplasia, and cleft lip/palate (EEC) syndrome patients carrying p63 mutations. Transcriptome analyses revealed distinct step-wise cell fate transitions during epidermal commitment; from multipotent simple epithelium to basal stratified epithelia, and ultimately to the mature epidermal fate. Differentiation defects of EEC iPSCs caused by mutant p63 occurred during the specification switch from the simple epithelium to the basal stratified epithelial fate. Single-cell transcriptome and pseudotime analyses identified signatures of embryonic epithelial-mesenchymal transition (EMT) associated with the deviated commitment route of EEC iPSCs. Repressing mesodermal activation reversed the EMT and enhanced epidermal commitment. Our findings demonstrate that p63 is required for specification of stratified epithelia, probably by repressing embryonic EMT during epidermal commitment. This study provides insights into disease mechanisms underlying stratified epithelial defects caused by p63 mutations and suggests potential therapeutic strategies for the disease.

**Significance statement:** Mutations in p63 cause several developmental disorders with defects of epithelial related organs and tissues including the epidermis. Our study is to dissect the unknown cellular and molecular pathomechanism. We utilized human induced pluripotent stem cells (iPSCs) derived from ectrodactyly, ectodermal dysplasia, and cleft lip/palate (EEC) syndrome patients carrying p63 mutations and studied transcriptome changes during differentiation of these cells to epidermal cells. Our analyses showed that the specification of the proper epithelial cell fate was affected by p63 EEC mutations, with an abnormal embryonic epithelial-mesenchymal transition (EMT). Repressing mesodermal activation reversed the EMT and enhanced epidermal commitment. This study provides insights into disease mechanisms associated with p63 mutations and suggests potential therapeutic strategies.

## Introduction

The transcription factor p63 is a key regulator in the development of stratified epithelia in many organs (1–3). Deletion of p63 in mice results in striking developmental defects or even complete absence of stratified epithelia in organs such as the epidermis, breast, prostate and bladder (2–5). In humans, heterozygous mutations in *TP63* encoding the p63 protein give rise to several autosomal dominant developmental disorders (6). These disorders manifest defects in tissues and organs where stratified epithelia (e.g. the skin and mammary glands) are present and where embryonic epithelial tissues are involved during development such as limb and orofacial structures (6). The human disease phenotypes resemble those in p63 knockout mice (2, 3), although in milder forms. One of these disorders is ectrodactyly, ectodermal dysplasia, and cleft lip/palate (EEC) syndrome (OMIM 604292) that is associated with mutations located in the p63 DNA-binding domain. EEC patients exhibit all the characteristic phenotypes of p63 mutation-associated diseases, namely, defects in the epidermis and epidermal related appendages, limb malformation and orofacial clefting. Hotspot mutations in the p63 DNA-binding domain affecting amino acids such as R204 and R304 have been reported to account for approximately 90% of the EEC cases (7). Consistent with the dominant inheritance pattern, these mutations have been proposed to have a dominant negative effect by abolishing p63 DNA binding (8, 9). How p63 EEC mutations affect development of stratified epithelia at the cellular and molecular level is however not yet understood.

The epidermis is probably the best studied model for p63 function. It has been established that p63 plays a pivotal role in epidermal keratinocytes (KCs), often termed as epidermal stem cells, and p63 orchestrates essential cellular programs including stem cell maintenance and proliferation, differentiation, adhesion and senescence (10–13). The function of p63 at the cellular level during epidermal development is however still under debate. Some studies reported that cells and tissues in p63 knockout mice lacked Krt5/Krt14 expression (2, 14), and showed retained Krt8 and Krt18 expression, concomitant with induced mesodermal genes (14, 15). These findings support a model in which p63 is essential for epidermal commitment, acting as a gatekeeper of the epithelial lineage (14). Consistent with this model, expression of p63 in mouse embryos is firstly induced in the surface ectoderm where Krt8 and Krt18 are expressed, and continuously increased during epidermal commitment (16). An alternative model is that p63 is not required for commitment towards the epidermal fate, as it was reported that a small number of Krt5 and Krt14 expressing cells was observed and that the morphology of cells and tissues was relatively normal in p63 knockout mice (5). One explanation for these opposite interpretations is that the cell fate determination in these studies was based on a limited panel of marker genes, or that observations were made from a small number of cells in a heterogeneous population. The analysis of the complete transcriptome, especially single-cell transcriptome that captures heterogeneous cell states (17–19) may resolve this issue. Furthermore, characterizing the cell states of cells and tissues carrying EEC mutations can provide insights into disease mechanisms of affected epidermis and other stratified epithelia.

Human induced pluripotent stem cells (iPSCs) have provided revolutionary tools for disease modeling and for testing potential therapeutic compounds using relevant patient material (20). In this study, we derived epidermal KCs from human iPSCs of healthy controls and EEC patients carrying a p63 mutation, R204W or R304W (21). Bulk and single-cell RNA-seq were performed to examine cell fate specification using an *in vitro* epidermal commitment model. The time course RNA-seq analysis during epidermal commitment of control iPSCs showed that the initial iPSCs entered an intermediate ‘multipotent’ phase before switching and committing towards the basal stratified epithelial fate followed by a maturation phase specific for the epidermal fate. Importantly, p63 mutant iPSCs failed to make the switch towards the epithelial fate, and showed enhanced non-epithelial cell identity. Epidermal commitment of both normal and p63 mutant iPSCs could be enhanced by compounds that can repress the alternative mesodermal fate.

## Results

### Generation of functional induced keratinocytes (iKCs) from iPSCs

To obtain robust epidermal commitment readouts, we used control iPSCs derived from two different individuals, generated from dermal fibroblasts donated by healthy individuals (for details of reprogramming, see Supplementary Material and Methods). One commonly used human embryonic stem cell line (ESC-H9) was also used in parallel as the control. Pluripotency and the differentiation capacity of the two iPSC lines were confirmed by expression of pluripotency marker genes OCT4 and SSEA4, and the *in vitro* differentiation assay for three germ layers (Supplementary Figure 1). All three lines together are termed as pluripotent stem cells (PSCs). To facilitate transcriptome studies, we established a complete feeder-free workflow (Figure 1A) by modifying a 30-day differentiation protocol of combining retinoic acid (RA) for inducing the ectodermal fate and bone morphogenetic protein 4 (BMP-4) for repressing the neuronal fate (22). Using this workflow, we directed the three control PSC lines towards the epidermal fate and examined the differentiation process at day 0, 4, 7, 15 and 30. The morphology of differentiating PSCs started to change on day 4 into a cobblestone shape that was similar to that of keratinocytes, and became more homogeneous at later stages during differentiation (Figure 1A). The basal epidermal markers p63 and KRT14 were induced weakly on day 4, became more visible on day 7, and were then progressively upregulated and consistently expressed in day-30 induced keratinocytes (iKCs) (Figure 1B and 1C). Epidermal commitment of these PSCs was also confirmed by marker gene expression at the mRNA level, namely downregulated pluripotency marker *OCT4*, firstly induced and subsequently decreased simple epithelial markers *KRT8* and *KRT18, and* upregulated *TP63* and *KRT14* (Figure 1D).

**Figure 1.**
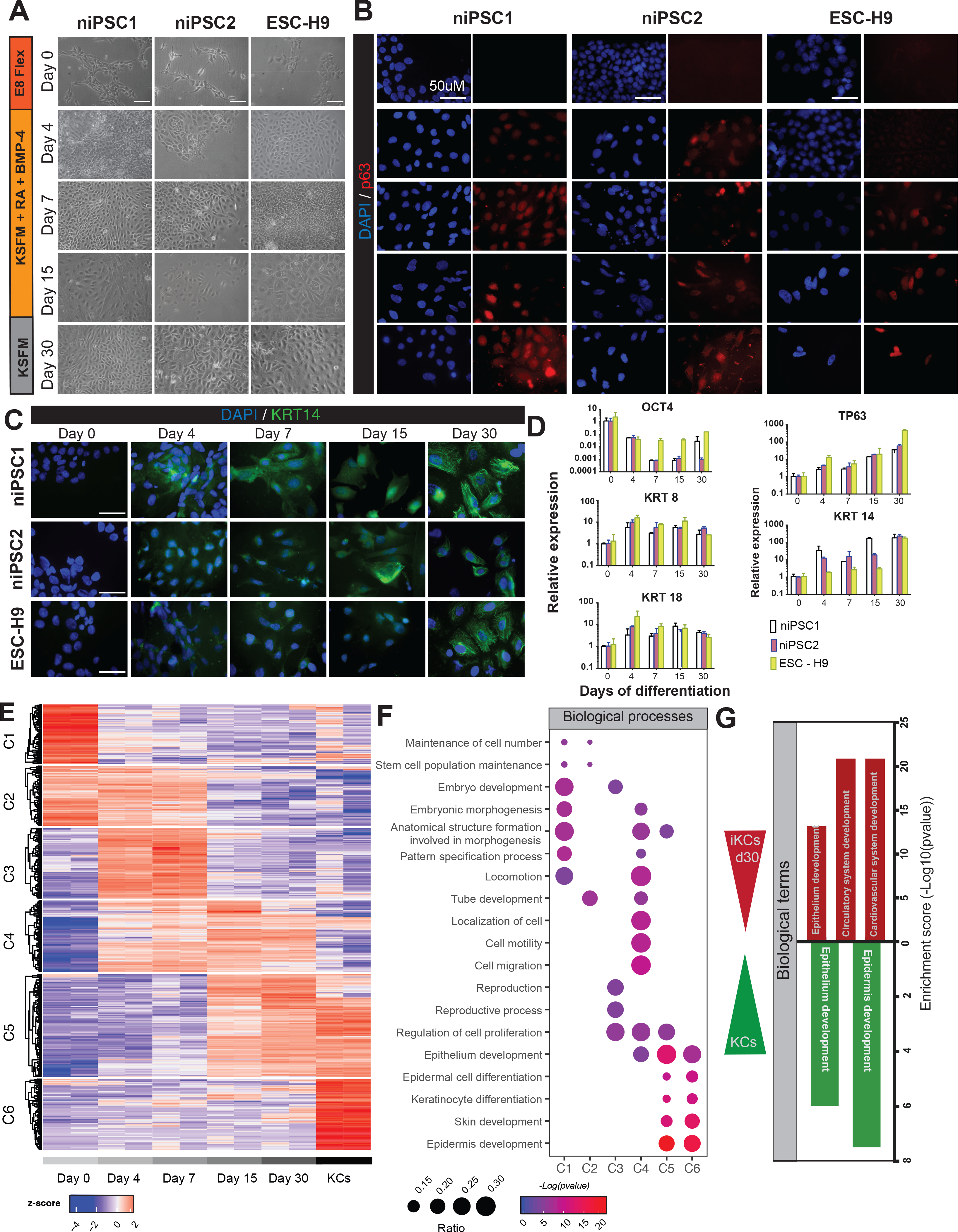
Generation and characterization of induced epidermal keratinocytes (iKCs) using human pluripotent stem cells (PSCs). (A) The schematic protocol for PSCs differentiation towards iKCs is perpendicularly presented to the bright field images of three different control PSCs cell lines (niPSC1, niPSCs2 and ESC-H9) that were induced for epidermal differentiation on day 0, 4, 7, 15 and 30 (Scale bars, 100 μm). (B)Immunofluorescence staining for p63 (red) on different days during differentiation (indicated in panel A). Cell nuclei were stained with DAPI (blue) (Scale bars, 50 μm). (C)Immunofluorescence staining for the epidermal marker *KRT14* (green) on different days during differentiation. Cell nuclei were stained with DAPI (blue) (Scale bars, 50 μm). (D)qRT-PCR analysis of the pluripotency marker *OCT4*, simple epithelium markers *KRT8* and *KRT18* and epidermal markers *TP63* and *KRT14* on different days during differentiation. Gene expression is expressed as 2^- ΔΔCt values over *GUSB* expression and undifferentiated day 0 samples (n=2 for group). Data are presented as mean ± SD, *p < 0.05, **p < 0.01, ***p < 0.001. (E)Heatmap of six clusters of differentially expressed (DE) genes in PSCs (niPSC1) during differentiation and in human primary keratinocytes (KCs). The colors in the heatmap indicate high (red) or low (blue) expression across the sample set. (F)Gene Ontology (GO) analysis showing biological process terms enriched for each cluster of DE genes. The gene ratio is indicated by the dot sizes and the significance by the color of the dot (Red: low pvalue: blue: high pvalue). (G)GO analysis showing biological process terms enriched for DE genes identified inthe pairwise comparison between day-30 iKCs and primary KCs. Terms enriched for DE genes that showed significantly higher expression in day-30 iKCs and those higher in primary KCs are indicated in red and green bars, respectively. The significance of the GO enrichment (*P* values) are indicated on the right axis.

Next we induced terminal differentiation/stratification of iKCs by including serum in the 2D culture medium, to assess whether the PSC-derived iKCs harbored stratification potential. Upon induction for 72 hours, induced expression of terminal differentiation markers, transglutaminase I (*TGM1*), involucrin (*IVL*) and cysteine protease inhibitor cystatin M/E (*CYSME*), was detected (Supplementary Figure 2A and 2B). These data demonstrated that these iKCs could be induced towards terminal differentiation. Taken together, iKCs derived from PSCs displayed both morphological and functional properties of epidermal basal cells and were similar to primary KCs.

To characterize the epidermal commitment process and molecular signatures of the derived iKCs, we performed time-course RNA-seq analyses of PSC cells during the 30-day differentiation process, and compared them with those of primary KCs (Supplementary Table 1). K-means clustering analysis of the top 500 differentially expressed (DE) genes during differentiation identified six distinct clusters (Figure 1E and 1F, Supplementary Table 2). Cluster 1 (e.g. *NANOG* and *OCT4*) and cluster 2 (*SALL4* and *SOX2*) were highly expressed in PSCs and contained genes involved in stem cell maintenance and early embryonic morphogenesis and development based on Gene Ontology (GO) analysis. Genes in cluster 3 such as *MSX1* and *CDX1* were transiently induced and subsequently switched off, and are involved in embryonic development (reproduction) as well as cell proliferation. Genes in cluster 4, e.g. *IGFBP3* and *GATA3*, were induced during early epidermal differentiation and stayed at a high level until differentiation day 30. Many of these genes seemed to play roles in cell migration and adhesion, embryonic morphogenesis and development of different organs such as epithelium, vasculature and skeletal system development. Importantly, cluster 5 included genes such as *KRT5, KRT17* and *TP63* that are involved in epithelial and epidermal development, desmosome assembly and extracellular matrix organization. These genes were lowly expressed in PSCs and during early differentiation, and significantly induced on day 15 and remained high on day 30. The expression levels of these genes were similar to their expression in KCs. Interestingly, cluster 6 also contained epidermal genes such as *KRT14, KRT1, IVL* and *FLG* that play roles in keratinocyte differentiation and barrier formation. However, genes of this cluster remained lowly expressed during the differentiation process, and even on day 30, their expression did not reach the levels in primary KCs. Gene expression of day-30 iKCs was similar but not identical to that of the primary KCs, which was evident from the expression of cluster 4 and 6 genes. These observations were also confirmed by Principal Component Analysis (PCA), indicating that PSCs went through distinct phases during epidermal commitment and there was apparent difference in the molecular signatures of day-30 iKCs and primary KCs (Supplementary Figure 2C and 2D, Supplementary Table 3).

To interrogate the difference between day-30 iKCs and epidermal KCs, we performed pair-wise comparison between these two conditions (Figure 1G; Supplementary Table 3). Gene that showed higher expression in day-30 iKCs were enriched for non-epithelial cell functions involved in circulatory and cardiovascular system development such as *ACTB* and *FBN1*, and for epithelial functions in other stratified epithelia such as *KRT18* and *KRT19*. In contrast, genes that were expressed lower in day-30 iKCs included those functioning in epidermal biology and adhesion, such as *KRT14, FGF2* and *LAMA3* (Figure 1G; Supplementary Table 4). These data suggest that, although morphologically (Figure 1A) and functionally (Supplementary Figure 2A, 2B) similar to primary KCs, day-30 iKCs had a distinct cell state as compared to primary KCs. They might be either immature KCs retaining embryonic epithelial signatures, or a mixture of basal epithelial cells that have the stratification potential.

### Differentiation defects of PSCs carrying p63 EEC mutations during epidermal commitment

Next, we examined the p63 function in epidermal specification by analyzing the effect of p63 EEC mutations during epidermal commitment. For this, we used two previously described iPSC lines derived from EEC patients who carry mutations in the p63 DNA binding domain, R204W or R304W (21), termed as EEC-iPSCs. The mutations in these iPSCs were confirmed by Sanger sequencing analyses (Supplementary Figure 3A). The morphology of control PSCs and EEC-iPSCs at the undifferentiated and early differentiation stages (day 0 and 4) was indistinguishable (Figure 2A; Figure 1A). Morphological differences between control PSCs and mutant iPSCs became evident from differentiation day 7. EEC-iPSCs were heterogeneous, lost the typical cobblestone shape and started to detach from culture plates. Few cells remained on culture plates on day 15 and did not survive till day 30 (Figure 2A). RT-qPCR analyses showed that *OCT4* expression decreased in all PSCs during differentiation, although the reduction was less pronounced in EEC-iPSCs (Figure 2B). Expression of simple epithelial genes *KRT8* and *KRT18* was not significantly different in EEC-iPSCs, as compared to that in control PSCs (Figure 2B and 2C), suggesting that the simple epithelium was not affected by EEC mutations. However, in contrast to the progressively increased expression level of epidermal markers p63 and KRT14 in control PSCs, the expression of these genes was significantly affected in EEC-iPSCs (Figures 2B, 2C and 2D). Expression of p63 seemed to be induced at day 4 but decreased on day 7 and especially on day 15 in EEC-iPSCs. KRT14 expression remained low at the mRNA level and was barely detectable at the protein level.

**Figure 2.**
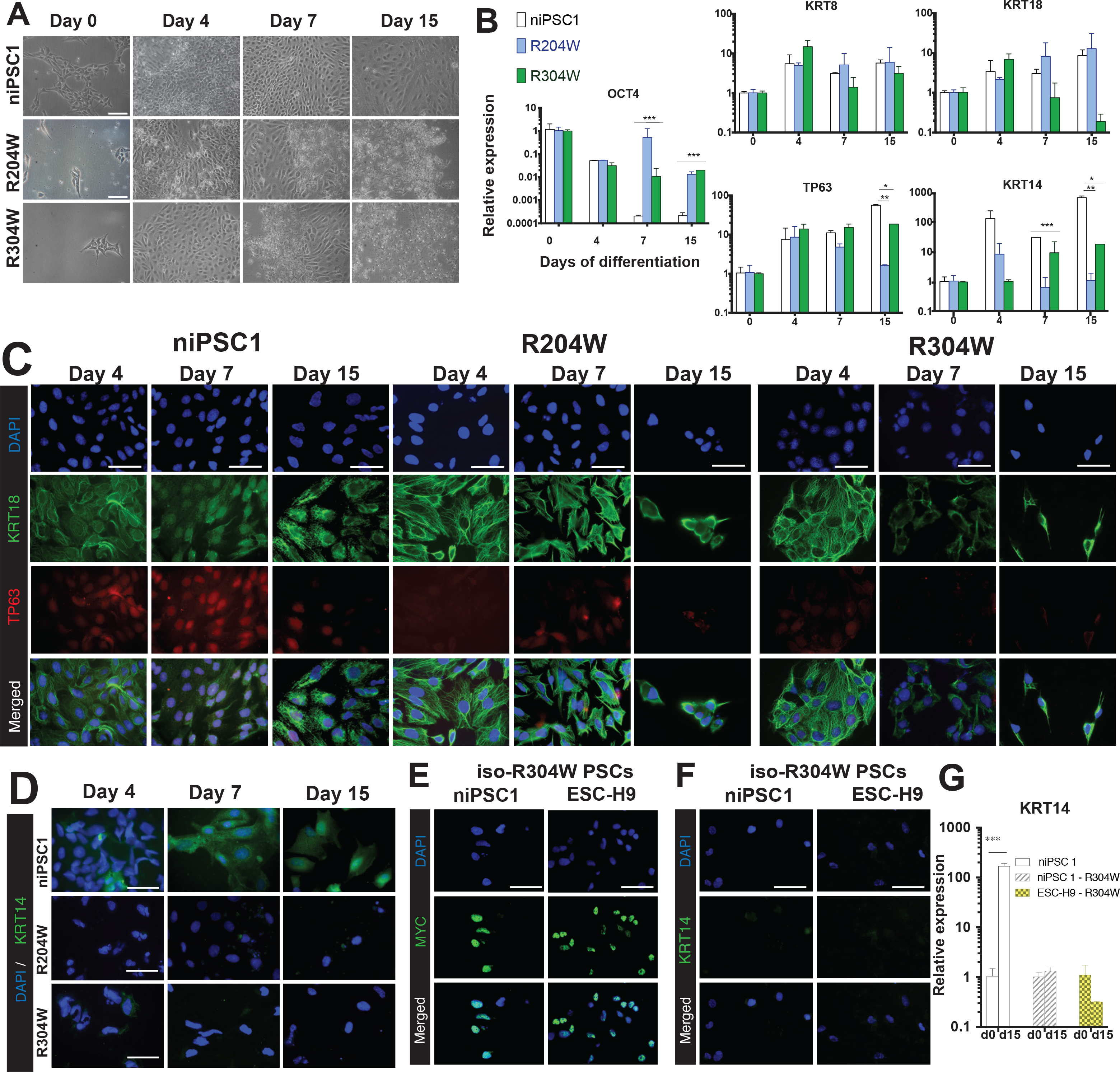
Differentiation defects of PSCs carrying p63 EEC mutations. (A)Bright field image of patient derived EEC-iPSCs, R204W and R304W,during epidermal differentiation till day 15. (B)qRT-PCR analysis of the pluripotency marker *OCT4*, simple epithelium markers *KRT8, KRT18* and epidermal markers *TP63* and *KRT14* for control niPSC 1 and EEC-iPSCs. Gene expression is expressed as 2^- ΔΔCt values over *GUSB* and undifferentiated day 0 samples for each cell line (n=2 for group). Student’s t-test was performed comparing values of differentiated cells against that of day 0, for each genotype. Data are presented as mean ± SD, *p < 0.05, **p < 0.01, ***p < 0.001. (C)Immunofluorescence staining for p63 (red), the simple epithelium marker KRT18 (green) and DAPI (blue) in niPSC1 and EEC-iPSCs on different days of differentiation (Scale bars, 50 μm). (D)Immunofluorescence staining for the epidermal marker KRT14 (green) and DAPI (blue) in niPSC1 and EEC-iPSCs on different days of differentiation (Scale bars, 50 μm). (E)Immunofluorescence staining for the MYC-tag (green) and DAPI (blue)in iso-R304W PSCs (niPSC1-R304W and ESC-H9-R304W) on day 15. (Scale bars, 50 μm). (F)Immunofluorescence staining for the epidermal marker KERATIN 15 (green) and DAPI (blue) in iso-R304W PSCs (niPSC1-R304W and ESC-H9-R304W) on day 15. (Scale bars, 50 μm). (G)qRT-PCR analysis of the epidermal marker *KRT14* for control niPSC 1, and iso-R304W PSCs. Gene expression is expressed as 2^- ΔΔCt values over *GUSB* and undifferentiated day 0 samples for each cell line (n=2 for group). Student’s t-test was performed comparing the committed cells against the day 0, for each genotype. Data are presented as mean ± SD, *px < 0.05, **p < 0.01, ***p < 0.001.

To corroborate that impaired commitment indeed results from p63 EEC mutations, we generated PSCs that contained the EEC R304W mutation on isogenic backgrounds of control PSCs. This was performed by introducing the mouse p63 R304W mutant into control iPSC1 and ESC-H9, termed as iso-R304W PSCs. Expression of the MYC-tagged p63 R304W mutant transgene was confirmed by immunostaining (Fig. 2E) and by sequencing of the cDNA derived from iso-R304W PSCs (Supplementary Figure 3B). During differentiation of these iso-R304W PSCs, heterogeneous cell morphology was observed (Supplementary Figure 3C), similar to the morphology detected during EEC-iPSC differentiation (Figure 2A). These differentiating iso-R304W PSCs seemed to survive slightly better than differentiating EEC-iPSCs․. Importantly, in contrast to the strong induction of epidermal marker *KRT14* in the isogenic control iPSC1 on differentiation day 15, no induction was observed in both iso-R304W PSCs (Figure 2D and 2G). RT-qPCR analyses showed similar marker gene expression during differentiation of iso-R304W PSCs, as compared to that in EEC iPSCs, such as downregulation of *OCT4* and a slight induction of *KRT8* and *KRT18* (Supplementary Figure 3D). The expression of *TP63* stayed unchanged (Supplementary Figure 3D), as the mouse p63 R304W mutant was expressed constitutively at all stages. These data unambiguously demonstrated that the expression and presence of the mutant p63 in PSCs abrogate the function of the normal p63 during epidermal commitment, which is consistent with the dominant negative model of p63 EEC mutations proposed previously (8, 23).

### Deregulated epithelial gene expression in differentiating PSCs carrying p63 EEC mutations

To characterize EEC-iPSC commitment at the molecular level, we performed a time course RNA-seq analysis of these cells during epidermal differentiation (Supplementary Table 1). K-means clustering analysis of the top 500 DE genes identified 7 clusters (Fig. 3A, Supplementary Table 5). Generally, down-regulated genes during differentiation, categorized into cluster 1 and 2, contained genes that are known to play roles in stem cell maintenance and the onset of differentiation, e.g *NANOG* in cluster 1 and *LIN28A* in cluster 2. The expression pattern of these two cluster genes between control PSCs and EEC-iPSC did not show apparent difference. Note that cluster 2 genes were not consistently regulated and no enriched biological function was detected based on GO annotation, and therefore this gene cluster probably represented the genetic background difference between different cell lines. Cluster 3 and 4 included genes that were initially induced and subsequently downregulated either during late differentiation (cluster 3) or in primary KCs (cluster 4). Expression of these two cluster of genes showed difference between control PSCs and EEC-iPSCs only at the early differentiation stage on day 4, and the difference diminished as differentiation proceeded. These genes including *MSX (MSX1)* and *WNT (WNT6)* mainly play roles in morphogenesis genes. Cluster 5 contained genes that were consistently expressed at a higher level in EEC-iPSCs, as compared to control PSCs, especially at later differentiation stages on day 7 and 15. These genes were strongly enriched for functions in immune and defense response, such as *IFNB1* and *IFIT1*. Cluster 6 included genes that were up-regulated during differentiation, and many are known for a role in epidermal development, e.g. *GJB3* and *KRT17*. For most of these genes, there was no obvious difference between control PSCs and EEC-iPSCs. Cluster 7 genes were strongly up-regulated in control PSC at late differentiation stages on day 15 and day 30, but remained consistently low in EEC-iPSCs. Genes in this cluster included *KRT5* and *DSG3* that have important roles in regulating epidermal development.

**Figure 3.**
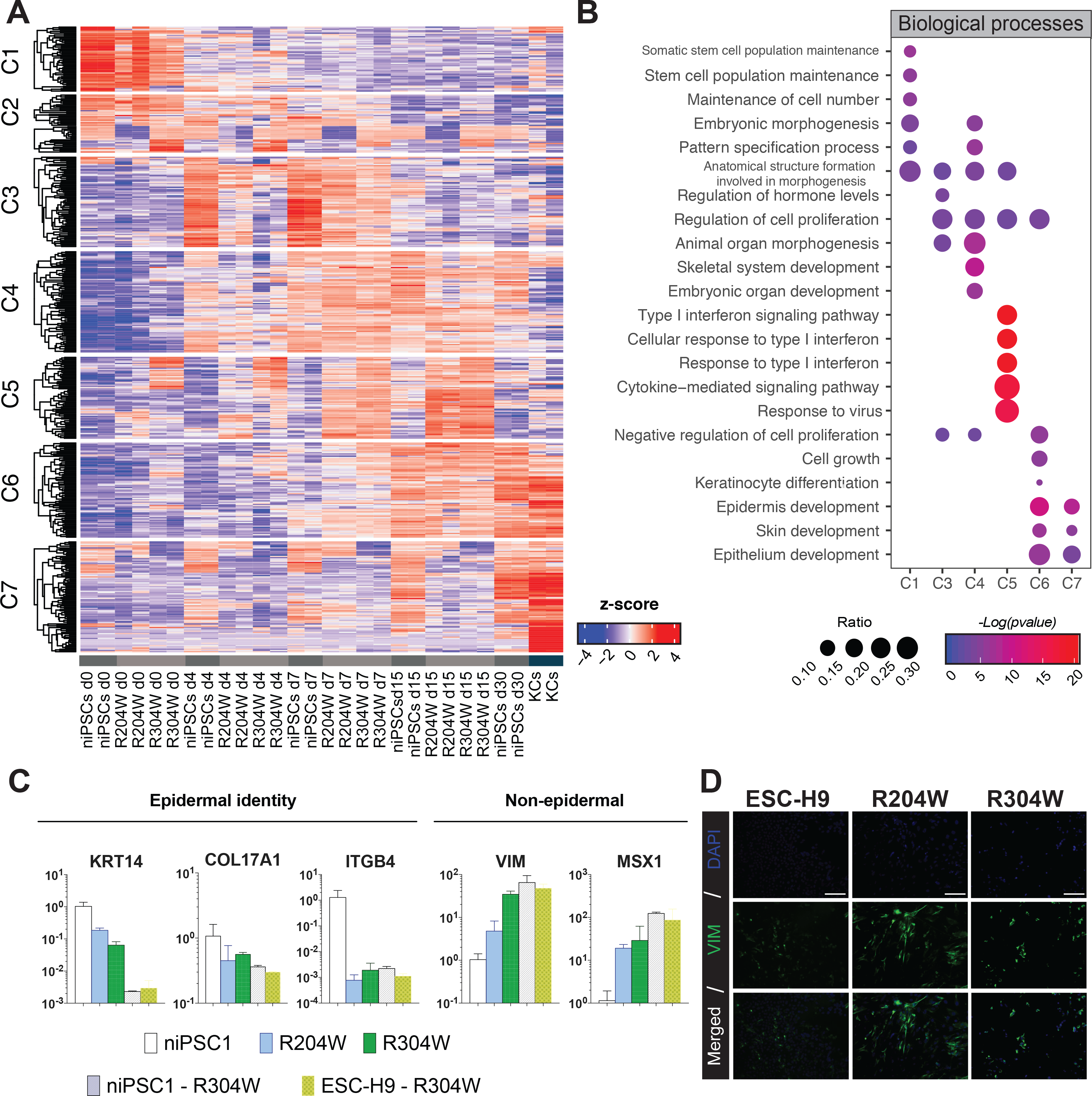
Deregulated epithelial gene expression in PSCs carrying p63 EEC mutations․. (A)Heatmap of eight clusters of differentially expressed (DE) genes in PSCs (niPSC1) and EEC-iPSCs during differentiation and in KCs. The colors in the heatmap indicate high (red) or low (blue) expression across the sample set. (B)Gene Ontology (GO) analysis showing biological process terms enriched for each cluster of DE genes. The gene ratio is indicated by the dot size and the significance by the color of the dot (Red: low pvalue: blue: high pvalue). (C)qRT-PCR analysis of epidermal genes KRT14, COL17A1 and ITGB4 and non-known epidermal genes VIM and MSX1 in control niPSCs, EEC-iPSCs and isoR304W-PSCs on differentiating day 15. Gene expression is expressed as 2^- ΔΔCt values over *GUSB* and niPSCs for each gene (n=2 for group). (D)Immunfluorescence of the mesodermal marker VIM in control ESC-H9 and EEC-iPSCs on day 15 (Scale bars, 100 μm).

The expression pattern of these seven clusters of genes was generally consistent with that in control PSCs during differentiation. Only two clusters among these seven clusters, cluster 5 and 7, showed a clear difference between control PSCs and EEC-iPSCs at later stages of differentiation (day 7 onwards) when p63 expression became readily detectable. This deregulated gene expression pattern, accompanied by differentiation defects of EEC-iPSCs observed at this stage, suggests that these two clusters of genes are likely directly affected by p63 EEC mutations. To further assess the gene expression difference between control PSCs and EEC-iPSCs, we looked for consistent DE genes that were common to both EEC mutations, R204W and R304W, on differentiation day 15 (Supplementary Figure 4A, Supplementary Table 6). Commonly downregulated genes (241) in EEC-iPSCs were related to epithelial and epidermal development (e.g. *KRT5, TP63* and *COL17A1*), whereas upregulated genes (296) in both EEC-iPSCs were enriched in non-epithelial functions such as immune response (Supplementary Figure 4B, Supplementary Table 6). We subsequently chose several typical epidermal and non-epidermal genes that were differentially regulated between control PSCs and EEC-iPSCs, and validated their expression in differentiating control PSCs, EEC-iPSCs and the isogenic iso-R304W PSCs by RT-qPCR analyses. Our analyses confirmed that epidermal genes *KRT14, COLA17A1* and *ITGB4* were consistently down-regulated in EEC-iPSCs and in isogenic iso-R304W PSCs, whereas non-epithelial genes *VIM* and *MSX1* were up-regulated, as compared to control PSCs (Figure 3C). The up-regulation of *VIM* in EEC-iPSCs was also confirmed at the protein level (Figure 3D). These data are in agreement with the previous study showing differentiation defects of EEC-iPSCs during epidermal and corneal epithelial commitment and upregulation of mesodermal genes, suggesting that the acquisition of the epithelial fate is impaired in EEC-iPSCs (21).

### Cell state heterogeneity during differentiation identified by single-cell RNA-seq

The next intriguing question is to identify the cell state of EEC-iPSCs that failed to commit to the epithelial fate. As differentiating PSCs may be heterogenous, we performed single-cell RNA-seq using a modified STRT-seq protocol (24, 25) to detect the heterogeneity of the cell population. In total, we analyzed the transcriptome of 1,250 single cells from control PSCs and EEC-iPSCs at different differentiation stages, as well as from primary KCs from a control and the same EEC patients carrying R204W and R304W. After filtering based on the data quality (Supplementary Figure 5), we obtained the transcriptome of 964 single cells. We detected on average 112k transcripts per sample that mapping to genes and 4,218 genes per cell. Following normalizations to get rid of batch effects (26), we detected 500 DE genes and performed PCA analysis (Figure 4A). Upon differentiation, DE gene expression of all PSCs initially moved diagonally along the major two axes PC1 (14.5%) and PC2 (11.4%). On day 15, DE genes reached the highest level along PC2, with those of EEC-iPSCs higher than those of control PSCs. Day-30 iKC DE genes moved down along PC2 but continued to move along PC1 towards primary KCs. Genes associated with PC1 showed an enrichment for functions in epidermal and epithelial development, whereas genes associated with PC2 were more enriched for migration, defense and extracellular matrix functions (Figure 4B, Supplementary Table 7). The triangle-shape transition during differentiation shown by PCA suggests that control PSCs went through an intermediated phase before committing to iKCs, and EEC-iPSC did not switch to the iKC fate, consistent with our bulk RNA-seq data. Interestingly, among the total number of 75 day-30 iKCs, six of them were close to primary KCs in the PCA analysis (Supplementary Figure 6A, black dots). When specifically examined, expression of several epidermal or non-epidermal marker genes in these six cells was more similar to their expression levels in primary KCs than in the bulk day-30 iKCs (Supplementary Figure 6C). This indicates that a small percentage (8%) of day-30 iKCs resembled primary KCs. Furthermore, t-Distributed Stochastic Neighbor Embedding (t-SNE) analysis showed that control PSCs and EEC-iPSCs before differentiation clustered together, and upon differentiation induction when p63 expression became detectable, differentiating control PSCs and EEC-iPSCs clustered into distinct groups (Figure 4C). These findings are in agreement with differentiation defects observed in differentiating EEC-iPSCs and with the data from the bulk RNA-seq analyses, suggesting that expression of mutant p63 gave rise to distinct cell states.

**Figure 4.**
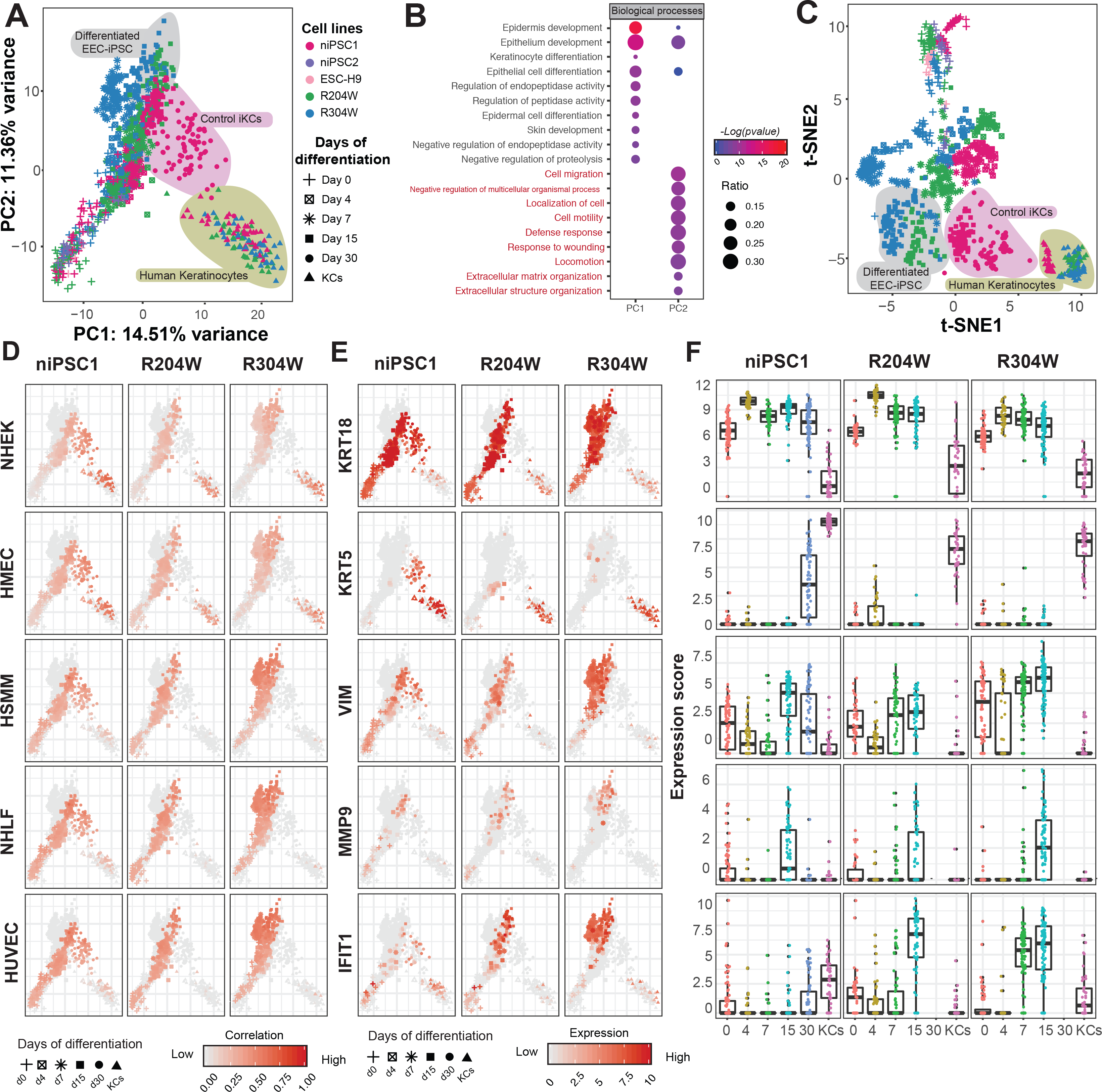
Cell state heterogeneity during differentiation identified by single-cell RNA-seq. (A)Principle Component Analysis (PCA) of the top 500 highly variable (HV) genes obtained from single-cell RNA-seq during epidermal differentiation of PSCs and primary keratinocytes. Colors represent cell lines; shapes represent differentiation days. (B)Gene Ontology (GO) analysis showing biological process terms enriched for PC1 and PC2 axes. The gene ratio is indicated by the dot sizes and the significance by the color of the dot (Red: low pvalue: blue: high pvalue). (C)t-Distributed Stochastic Neighbor Embedding (t-SNE) analysis of the top 500 highly variable (HV) genes obtained from single-cell RNA-seq during epidermal differentiation of PSCs and primary keratinocytes. (D) Correlation of single cell transcriptome against bulk RNA-seq data of cell types from diverse lineages shown in PCA (A). NHEK, normal human epidermal keratinocytes, HMEC, human mammary epithelial cells, HSMM, human skeletal muscle myoblasts, NHLF, normal human lung fibroblasts, HUVEC, human umbilical vein endothelial cells. Correlation coefficient is indicated by the color. Red: high correlation; gray: low correlation and other cell types. Marker gene expression levels (Log(TPM+1)) in single-cell transcriptome shownin PCA (A). Simple epithelium marker, KRT18; epidermal marker, KRT5; mesodermal marker, VIM, MMP9 and IFIT1 Red: high correlation; gray: low correlation and other cell types. (E) Marker gene expression levels (Log(TPM+1)) in single-cell transcriptome shown by bar plots. Marker genes are labeled in panel (E).

To identify the cell fates during different stages of differentiation, we deconvoluted bulk RNA-seq data of cells derived from different embryonic origins (GEO: GSE101661) (27) using our single-cell transcriptome. As expected, the starting PSCs including both control PSCs and EEC-iPSCs showed high correlations with ESC-H9 bulk RNA-seq data, and the correlations decreased when PSCs were induced for differentiation (Supplementary Figure 6D). In addition, primary control KCs, EEC-KCs and day-30 control iKCs of showed low correlation to human ESC-H9 data, but highest correlations with the two types of epithelial cells, Normal Human Epidermal Keratinocytes (NHEK) and Human Mammary Epithelial Cells (HMEC) (Figure 4D). These high correlations were also consistent with the high expression level of *KRT5* and *ITGA6* in these cell types, despite the heterogeneous expression of these genes, especially in day-30 iKCs (Figure 4E and 4F; Supplementary Figure 7A and 7B). Interestingly, day-30 control iKCs showed relatively high correlation to non-epithelial cell types, Human Skeletal Muscle Myoblasts (HSMM), Normal Human Lung Fibroblasts (NHLF) and Human Umbilical Vein Endothelial Cells (HUVEC), and ESC-derived cells such as non-neural ectodermal (NNE) and neural crest cells (NCS). This was distinct from primary KC that showed low or no correlation to these non-epithelial cells (Figure 4D, Supplementary Figure 6C). Consistent to the correlations, control day-30 iKCs showed lower expression of *KRT14* and *DSG1* but higher expression of *KRT18* and *KRT19*, as compared to primary KCs (Figure 4E and 4F; Supplementary Figure 7). *KRT18* and *KRT19* are known to be often highly expressed in other epithelial cells such as in mammary glands or oroepithelial cells (28). Furthermore, it is noticeable that differentiating control PSCs on day 15 showed the highest correlation with embryonic epithelial and non-epithelial cells and their corresponding markers (Figure 4D, 4E, 4F, Supplementary Figure 6D and 7). In agreement, the high expression of the non-epithelial marker *VIM* was detected at this stage (Figure 4D; Supplementary Figure 6C). These data indicate that a multipotent cell state is established at early stages during normal PSC differentiation, before a switch towards the epidermal fate. The multipotent state however is partially retained in day-30 iKCs that probably represent a basal stratified epithelia fate.

When examining the cell states of differentiating EEC-iPSCs on day 15, the difference of cell state correlation between epithelial cell types (NHEK and NMEC) and EEC-iPSCs seemed not to be obvious, neither was the expression difference between simple epithelial marker *KRT18* or epithelial markers *KRT5* and *KRT14* (Figure 4D, 4E, 4F; Supplementary Figure 6C, 7A, 7B). The most apparent difference was detected in the higher correlations of EEC-iPSCs with non-epithelial cells, HSMM, NHLF, HUVEC and ESC-derived other cell types, as compared to control PSCs on day 15 (Figure 4D; Supplementary Figure 6C). The correlations were consistent with higher expression of non-epithelial genes such as *VIM*, *MMP* and *IFIT* genes in EEC-iPSCs, especially in R304W-iPSCs (Figure 4E and 4F; Supplementary Figure 7A and 7B). These findings indicate that EEC-iPSC exhibited an enhanced non-epithelial cell state, arguing that they retained the multipotent cell state and that differentiation defects caused by p63 EEC mutations occurred during the switch to the epidermal fate.

In summary, the single-cell transcriptome data demonstrated the cell state heterogeneity during *in vitro* PSC epidermal commitment. Collectively, our temporal transcriptome analyses indicate that the normal epidermal commitment process consists of several connected yet distinct phases: the phase of establishing multipotency where PSCs exit the pluripotent state upon differentiation induction and acquire a transient multipotent fate; the phase of specification of basal stratified epithelia where differentiating PSCs switch towards the embryonic pre-mature epidermal fate or a mixture of basal epithelial fate; and the maturation phase where cells are specified to epidermal KCs. Furthermore, differentiating PSCs carrying EEC p63 mutations retained non-epithelial cell identity, suggesting that EEC p63 mutations disturbed the switch from the multipotent state to the basal stratified epithelial fate, and p63 expression is not sufficient for epidermal maturation.

### Deviated commitment route with enhanced non-epithelial cell fates in EEC-iPSC differentiation

To identify the timing of the affected cell state switch and the potential alternative differentiation route of EEC-iPSCs, we examined differentiation trajectories of PSCs towards KCs using pseudotime analysis Monocle (17). When cells were ordered according to gene expression pseudotime, control PSCs exhibited a switch at day 7 and day 15, and then progressed on the path towards primary KCs, even though most day-30 control iPSCs did not reach the primary KC fate, suggesting that day-30 control iPSCs retain some embryonic multipotent signatures (Figure 5A). In contrast, differentiating EEC-iPSCs diverted towards other directions. The pseudotime path of R304W-iPSCs showed more sub-branches than that of R204W-iPSCs, suggesting a more severe differentiation defects (Figure 5A). When only the transcriptomes of differentiating PSCs were analysed without including the primary KC profiles, the bifurcation that occurred on day 7 and day 15 between control PSCs and EEC-iPSCs became more apparent (Figure 5B). The pseudotime path of all differentiating R304W-iPSCs and the majority of R204W-iPSCs took an alternative direction (branch 2 in Figure 5B) to the pseudotime path of day-15 and day-30 control PSCs. According to the pseudotime ordering, differentially regulated genes were clustered into six clusters, and cluster 1 and 2 genes corresponded to the bifurcation branches 1 and 2, respectively (Figure 5C). GO analyses of cluster 1 genes showed that these genes were associated with epidermal development and keratinocyte differentiation; whereas that of cluster 2 genes showed an enrichment of genes involved in immune response (Figure 5D). In agreement, correlation analyses to bulk RNA-seq data of other cell types showed that branch 1 and branch 2 genes had higher correlations to NHEK cells and to non-epithelial HSMM cells, respectively (Figure 5E; Supplementary Table 8). Genes in other clusters of the psudotime analysis corresponded to genes that were expressed earlier during differentiation (Figure 5B, Figure 5C). These data demonstrated that the differentiation route of EEC-iPSCs deviated from the normal commitment route towards the epithelial fate. The deviation occurred at the multipotent state on differentiation day 7 and 15 when p63 expression was detectable, prior to epidermal maturation.

**Figure 5.**
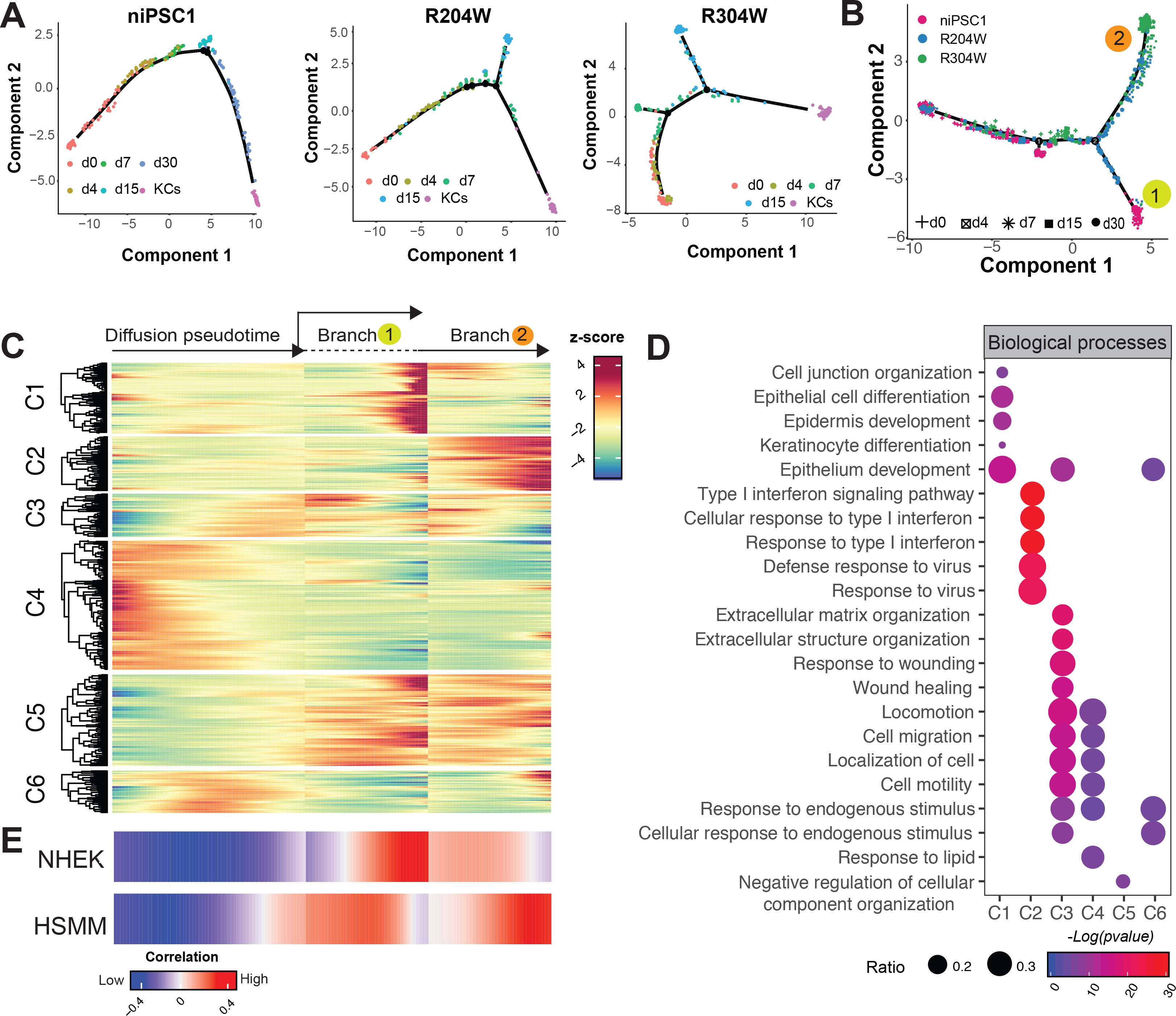
Deviated commitment route with enhanced non-epithelial cell fates in EEC-iPSC differentiation. (A)Differentiation gene expression trajectories of control niPSC1 and EEC-iPSCs during differentiation and primary KC using Pseudotime (Monocles) analysis. Differentiation days are indicated by colors. (B)Differentiation gene expression trajectories of control niPSC1 (pink) and EEC-iPSCs, R204W-iPSCs (blue), R304-iPSCs (green) during differentiation using Pseudotime (Monocles) analysis, without primary KCs. Differentiation days are indicated by shapes.(C) Heatmap of DE genes detected by Pseudotime analysis based on their Z-scores (indicated by the color). (D)GO analysis showing biological process terms enriched for HV genes in each cluster. The gene ratio is indicated by the dot size and the significance by the color of the dot (Red: low pvalue: blue: high pvalue). (D)Correlation of single cell transcriptome against bulk RNA-seq data of NHEK (normal human epidermal keratinocytes) and HSMM (human skeletal muscle myoblasts). Correlation coefficient is indicated by the color.

### Epidermal differentiation enhanced by compounds that repress mesodermal differentiation

As differentiating EEC-iPSCs had higher expression of immune response genes and a higher correlation with the HSMM cell identity, we reasoned that these differentiating EEC-iPSCs exhibited an enhanced mesodermal identity, as both the immune system and skeletal muscles derive from the mesoderm. As it has been reported that p63 can repress epithelial-mesenchymal transition (EMT) in cancers (29, 30), we hypothesized that normal p63 may have similar function in inhibiting the embryonic EMT during development, and EEC mutations affect the EMT repression controlled by p63. To assess whether the mesodermal activation is causal to the commitment defects of these EEC iPSC, three compounds that were shown to repress the mesodermal lineage differentiation, Heparin, valproic acid (VA) and Suramin (31–35) were tested. As our day-30 iKCs seemed to retain some multipotent signatures, repressing mesodermal differentiation may also facilitate epidermal maturation. These compounds were added to the differentiation medium from day 7 onwards, given that at this time point on cells entered a multipotent state, and p63 started to be expressed and exerted its role for epidermal commitment (Figure 6A).

**Figure 6.**
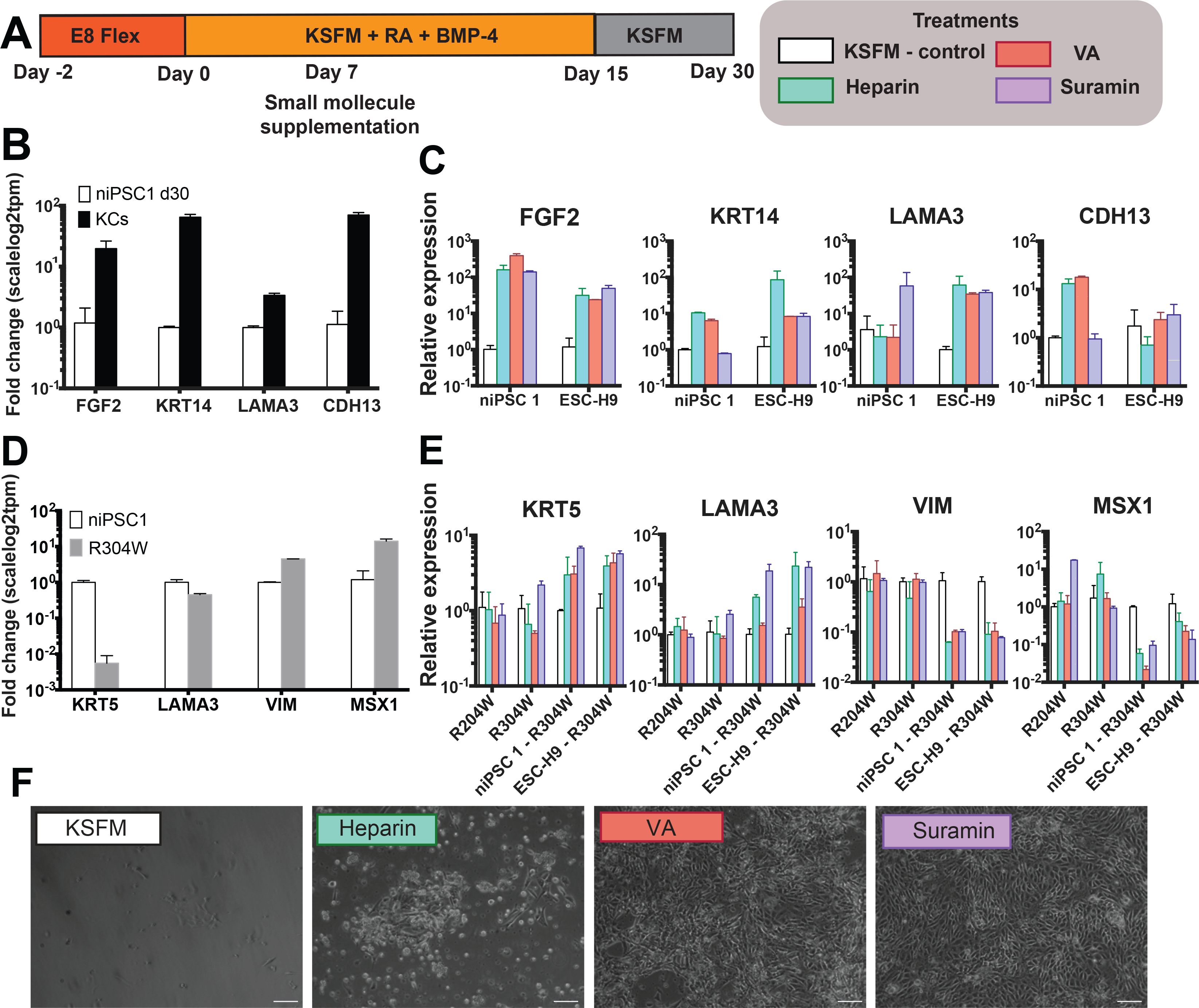
Epidermal differentiation enhanced by compounds repressing the mesodermal fate. (A) Scheme of suppplementation of small molecules, Heparin, Valproic acid (VA) or Suramin, during epidermal differentiation. (B) Fold change difference (scalelog2tpm) of gene expression between day-30 iKCs and control KCs, detected by bulk RNA-seq. (C) qRT-PCR analysis of in day-30 control iKCs supplemented treated with small molecules. Gene expression is expressed as 2^- ΔΔCt values over *GUSB* and day-30 iKCs without treatment (n=2). (D)Fold change difference (scalelog2tpm) of gene expression between niPSC1 and R304W on day 15, detected by bulk RNA-seq. (E) qRT-PCR analysis in day-15 R304W-PSCs supplemented treated with small molecules. Gene expression is expressed as 2^- ΔΔCt values over *GUSB* and day-15 R304W-PSCs without treatment (n=2 for group). (F) Bright field images of day-15 R304W-iPSCs without (KSFM) and with small molecules treatment (Heparin, VA, Suramin).

Upon treatment by the compounds, expression of several epidermal genes, *KRT14, LAMA3* and *CDH13*, whose level was lower in day-30 iKCs than that in primary KC (Figure 6B) was significantly elevated (Figure 6C). The effect of Heparin and VA seemed more consistent than that of Suramin. In addition, FGF2 that was lowly expressed in day-30 iKCs was also significantly induced upon the treatment by all three compounds (Figure 6B and 6C). To test effects of these compounds on differentiation of PSCs carrying p63 EEC mutations, we chose to use the R304W mutant, as this R304W-iPSCs gave the most severe defects during differentiation and an additional iso-R304W PSC line was available to be tested in parallel. Without treatment, differentiating R304W-iPSCs and iso-R304W PSC suffered from severe cell death in the normal keratinocyte medium KSFM at differentiation day 15 (Figure 2A and 6F). Upon the treatment, cells survived beyond day 15, especially in the Suramin-containing medium, differentiating PSCs had a cobblestone morphology that was similar to primary KCs on day 15 (Figure 6F). Gene expression analyses by RT-qPCR showed that the treatment by these compounds rescued deregulated gene expression in differentiating R304W-PSCs and gave rise to upregulated epidermal genes such as *KRT5* and *LAMA3* and downregulated mesenchymal genes such as *VIM* and *MSX1* (Figure 6D and 6E). Taken together, we showed that repressing the mesodermal activation could enhance epidermal commitment, suggesting that the embryonic EMT is a reversible process.

## Discussion

The role of p63 in controlling the integrity of different stratified epithelial tissues, especially how p63 mutations associated with developmental disorders affect these tissues, remains largely unexplored. In this study, we used epidermal commitment as the model to study cellular and molecular signatures of cell fate transitions from the pluripotent cell fate to stratified epithelia and how this process is affected by p63 mutations associated with EEC syndrome. Using bulk and single-cell transcriptome analyses, we showed that PSCs commitment towards the epidermal fate includes several connected yet distinguishable phases: 1) PSC commitment towards the multipotent simple epithelial fate; 2) the switch from the simple epithelium to the basal stratified epithelial fate; and 3) maturation of the epidermal fate. Single-cell analyses identified that differentiation defects associated with p63 EEC mutations occurred during the switch from the multipotent simple epithelium to the basal stratified epithelial fate, prior to epidermal maturation, and expression of p63 was not sufficient for epidermal specification and maturation. As the differentiation route of EEC PSCs exhibited enhanced mesodermal signatures, we tested compounds that repress the mesodermal differentiation and showed that repressing mesodermal activation could enhance epidermal commitment.

Stratified epithelia are multi-layered structures found in many organs as the outer surface such as the epidermis or lining of internal organs such as the bladder. Although they may be different from their biological functions and from their embryonic origins, derived from distinct germ layers, stratified epithelia share similarities in their transcriptional programs that dictate differentiation and development of these tissues (4, 5, 36). For example, high expression of *KRT5* and *KRT14* is found in the basal cells of these epithelia. Therefore, simply relying on cell and tissue morphology and expression of a limited number of mark genes to study cell fate specification and commitment is probably not sufficient for distinguishing similar cell states. In this study, we performed a comprehensive transcriptome analysis that is mandatory to reveal the cell identity at the molecular level. During epidermal commitment, although PSCs started to change morphology to a cobblestone shape that was similar to primary KCs already on differentiation day 4, the molecular signature represented a simple epithelial signature. The day-30 iKCs showed several molecular hallmarks of epidermal cells, with upregulated *KRT5* and *KRT14* and downregulate *KRT8* and *KRT18* (Figure 1A-1D), and could even be further induced towards the terminal stratification (Supplementary Figure 2A and 2B). However, our transcriptome analyses showed that these day-30 iKCs did not have the same molecular signature as the primary KCs, with quantitatively lower *KRT5* and *KRT14* expression and higher *KRT8* and *KRT18* expression (Figure 1E-1G). This indicates that day-30 iKCs may retain some embryonic properties. Alternatively, they may represent a mixture of basal epithelial cells that can stratify, as *KRT8* and *KRT18* are expressed in some stratified epithelial cells such as those in mammary glands (37). In line with these interpretations, day-30 iKCs shared high similarity to other epithelial (HMEC) or embryonically derived (NCS and NNE) cell types (Figure 4D and Supplementary Figure 6C).

Interestingly, our single-cell transcriptome analysis showed that 6 out of 79 cells (^~^8%) in day-30 iKCs had similar molecular signatures to primary KCs (Supplementary Figure 6A). This suggests that, with a careful and prolonged selection of the day-30 iKCs, e.g. enrichment for marker gene expression, it is possible to obtain a reasonable population of iKCs that resemble the molecular signature of primary KCs and can subsequently be cultured for generation of a 3D organotypic skin equivalent (38, 39). The low percentage of iKCs that faithfully resemble primary KCs, however, emphasizes the necessity of improving the epidermal differentiation protocol. Our effort in testing compounds that inhibit mesodermal activation gave rise to the promising lead that should be further evaluated. Our findings also highlight the importance of using comprehensive (single-cell) transcriptome analysis to identify molecular signatures and proper cell states.

Previous studies reported that p63 is essential for proper epidermal commitment (14, 40). Recent exciting new findings demonstrated the role of p63 in controlling the chromatin, especially the enhancer landscape in KCs (41, 42). Our study dissected the epidermal commitment and maturation process into further details, showing that connected yet distinct phases are involved in the process. Proper p63 expression and function are required for the switch from the multipotent state to the basal stratified epithelial fate, but not sufficient for epidermal maturation. p63 EEC mutations disturbed this switch, prior to the maturation of the epidermal cells. The detrimental function of p63 during the switch for specification of the basal stratified epithelial fate is consistent with defects in all stratified epithelia observed in p63 deficient mouse models (2, 3, 43) and in human diseases associated with p63 mutations (44). In line with previous observations of upregulated mesodermal gene expression in p63 deficient models (15, 45, 46), we showed that the mesodermal cell identity was enhanced in PSCs carrying p63 EEC mutations during differentiation. However, rather than simply being blocked between the ectodermal and fully committed epidermal stage, suggested by p63 deficient mouse studies, our single-cell RNA-seq and psedotime analyses showed that the differentiating mutant PSCs deviated from the normal route of epidermal commitment to a more mesodermal directed cell identity (Figure 5). A role of p63 in repressing EMT has previously been reported for epithelial related cancers, such as squamous cell carcinoma and breast cancer (29, 30). This intrigues us to propose that p63 plays a similar role in repressing the embryonic EMT during development of stratified epithelia, and p63 EEC mutations disrupt the normal function of p63 in repressing EMT. Our experiments using small molecule compounds showed that inhibition of mesodermal activation could reverse the EMT and enhance epidermal commitment. Given that p63 is normally induced after the multipotent state is established (differentiation day 4), differentiation defects observed in PSCs carrying p63 EEC mutations at later stages (on day 15) was likely due to the impaired p63 DNA binding to its target genes. The role of p63 in upregulating epithelial and epidermal genes directly has been well studied (11, 12, 47, 48). It would be of great interest to investigate whether p63 can repress the mesodermal genes by binding directly to these gene promoters or enhancers or via activating genes that can repress the mesodermal differentiation.

All three compounds tested for epidermal commitment in this study, Heparin, Valproic acid (VA) and Suramin, are known to repress the mesodermal lineage or interfere with EMT (34, 35, 36–38). Heparin and VA seemed to be more effective in improving epidermal KC maturation, whereas Suramin was more efficient in rescuing differentiation defects of EEC-PSCs. These differences may probably result from different working mechanisms of these compounds. Heparin can modulate FGF and EFG signaling (49, 50). VA is a histone deacetylase inhibitor that can upregulate H3 actylation and repress EMT (51). Suramin can inhibit mesoderm formation (52) and is well known to have antiviral effects (53, 54). For example, it has been reported that Heparin has strong binding to FGF2 at the basal membrane of the epidermis (49). FGF2 expression was lower in day-30 iKCs than in primary KCs (Figure 6B). In day-30 iKCs treated with the three compounds, FGF2 expressed was increased, with concomitant enhanced expression of typical epidermal markers such as *KRT14, LAMA3* and *CDH13* (Figure 6C). It is plausible that Heparin cooperates with upregulated FGF during epidermal maturation. However, the action of these compounds in enhancing epidermal commitment by repressing the mesodermal identity would be of importance to further investigate. Nevertheless, our rationalized compound testing provides promising future directions for improving epidermal commitment protocols and therapeutic development for diseases associated with p63 mutations, not only for the phenotype of the epidermis but also of other stratified epithelia.

It should be noted that differentiation defects observed in our *in vitro* human PSC epidermal differentiation model was more severe than in EEC patient skin, as the skin phenotype of EEC patients is rather mild (6) and basal epidermal cells are present. This is probably due to the *in vivo* heterogeneous cell and tissue environment where other signals and pathways may compensate, e.g. those from the dermis that are probably independent on p63 (55, 56). Nevertheless, our simplified *in vitro* differentiation assay sheds lights on the direct function of p63 and its regulatory gene network that is relevant to cell states and function. Our study here provides novel insights into the master regulatory function of p63 in early epidermal commitment and mechanisms underlying defects of stratified epithelia in disorders associated with p63 mutations.

## Material and Methods

Detailed descriptions of methods are presented in Supplementary Information (SI).

### Control iPSCs generation and characterization

Dermal fibroblasts of two health control individuals were reprogrammed into iPSCs by lentiviral transduction of human *OCT4, SOX2, KLF4* and *cMYC* factors using the hOKSMco-idTomFRT plasmid (57). Human iPSC generated in this study were characterized for their differentiation potential into the three germ layers using Human Three Germ Layer Kit (R&D systems),according to the manufactures’ instructions with minor modifications. iPSCs carrying *TP63* mutations were described and characterized previously (21).

### Differentiation of human PSCs into iKCs

PSCs were seeded as single cells before the initiation of differentiation. One or two days after seeding, differentiation was induced by incubating cells in keratinocyte induction medium (keratinocyte serum free medium (KSFM, GIBCO) + 10 ng/mL Bone Morphogenetic Protein-4 (BMP-4, PromoKine) + 0.3 mg/mL retinoic acid (RA, Sigma-Aldrich). After 7 days cells were dissociated and seeded in Geltrex coated plates with induction medium, and kept in induction medium until day 15. From day 15 to day 30 cells were cultured in KSFM only. In the experiments for testing small molecule compounds, cells were cultured from day 7 in induction medium supplemented with either 1mg/mL Heparin Sodium Salt (Heparin, Sigma-Aldrich), 0.25uM Valproic Acid (VA, Sigma-Aldrich) or 100uM Suramin (Santa Cruz). After day 15 cells were cultured with KSFM with small molecules. The medium was refreshed every other day and dishes were incubated at 37°C and 5% CO_2_. Stratification of iKCs were induced in the basal keratinocyte medium KSFM with 10% of Fetal Bovine Serum (FBS) (GIBCO), when cells were at 60% confluency. Coverslips and RNA were collected prior induction, 0 hours as well 48 and 72 hours after induction.

### Generation of lentiviral constructs, virus production and transduction

Lentiviral constructs were made through MultiSite Gateway 3-Fragment Recombination Reaction (Invitrogen). pEntry221-EFL-promoter and pEntry201-p63mutantR304W were recombined into the 2K7bsd lentivector (58) using LR Clonase II Plus Enzyme Mix (Invitrogen). For transduction, cells were cultured in Essential 8 medium and 50μL of p63 R304W mutant virus was added. Selection of infected cells were made in presence of 6 μg/ml Blasticidin S HCl (GIBCO) for at least 2 weeks before starting differentiation.

### RNA extraction, qRT-PCR and statistical analysis

Total RNA was isolated at day 0 or 4, 7, 15 (WT and EEC-iPSC) and 30 (for WT only) days after the induction of epidermal commitment. Cell aggregates were lysed on plate using 350 μL RA1 (NucleoSpin^®^ RNA kit (Macherey-Nagel) buffer with 3,5 μL β-mercaptoethanol (Sigma-Aldrich) and snap-frozen in liquid nitrogen. Total RNA was extracted with NucleoSpin^®^ RNA kit (Macherey-Nagel) and converted into 100-500ng cDNA using the iScript cDNA synthesis kit (Bio-Rad). RT-PCR was performed with 10–100 ng of cDNA template in a 25 μl total reaction volume (12.5 μl iQ SYBR Green Supermix (2×) (Bio-Rad), 0.5 mM of each gene-specific primer, H_2_O up to volume). Glucuronidase Beta (*GusB*) was used as housekeeping gene to normalize cDNA levels. The significance of the qRT-PCR results was determined by Student’s t-test and represented as mean ± SEM followed by a Bonferroni post-hoc test with the software Graphpad Prism 6.0. P-values below 0.05 were considered significant and are indicated in the figures.

### Immunostaining

Cells were seeded in Geltrex (GIBCO) coated coverslips. At different days of differentiation coverslips were harvested. They were fixed in 4% paraformaldehyde/1 × PBS for 15 min at room temperature or in methanol (Sigma-Aldrich) for 20min at −20°C. Slides were mounted in VECTASHIELD Antifade Mounting Medium with DAPI (Vectorlabs). Images were acquired in a Leica DM Fluorescence Microscope.

### Bulk RNA Sequencing Library Preparation

The NuGEN Ovation RNA-Seq (version 2, NuGEN) protocol was carried out on 100 ng of initial RNA. RNA-seq libraries were amplified using 10 cycles. The purified PCR products were then quantified using an Agilent bioanalyzer (DNA-1000 kit). We selected 300bp fragments using E-Gel™ SizeSelect™ II Agarose Gels, 2% (E-Gel, GIBCO). DNA fragments from the complete and size-selected library were detected and confirmed, respectively, using an Agilent bioanalyzer (DNA-1000 or High sensitivity kit). Ready to sequence RNA-seq libraries (size-selected) were tested for the expression of lineage-specific genes and quantified with a KAPA library quantification kit (KAPA Byosystems). All sample libraries were prepared were sequenced on Illumina HiSeq 2500 (Illumina, San Diego, CA, USA) for paired-end reads of 75×75bp.

### Data Analysis

Sequencing data were processed by standard methods for mapping (to the human hg19 using hisat2, version 2.1.0), differential gene expression (DESeq2, version 1.18.1, with adjusted p-value < 0.05 as the cutoff) and GO terms analyes (using Toppgene (https://toppgene.cchmc.org/), only Biological Process terms were kept for downstream analyses. All PCA and tSNE scater plots and GO-term results were generated with the ggplot2 (version 2.2.1) package. All the heatmaps were generated using the ComplexHeatmap version 1.17.1) package with centered and scaled log_2_(TPM+1) score.

### Single cell RNA sample preparation

A modified STRT-seq protocol (24) was for the generation of single-cell transcriptome profiles. The main steps are described in SI.

### Single cell RNA-seq analysis

For each single cell sample, read counts representing gene expression levels were calculated using HTSeq (version 0.9.1) with default parameters. Only protein coding genes and ERCC gene counts information was kept and continued for downstream analyses. Expression levels in all samples were normalized with the Scatter (version 1.5.0) R/Bioconductor package by log_2_(TPM+1)

For all 1,250 sequenced single cells, the low quality samples were filtered with five metrics: mapping rate, library size, gene number, percentage of counts mapping to mitochondrial genes and to ERCC transcripts. The library size and gene number threshold were defined by the median absolute deviation (MAD) of above 3, the percentage of counts mapping to mitochondrial genes and to ERCC transcripts was defined by the MAD of below 3. In total, 964 single cell samples were kept for downstream analyses. We used RUVSeq R package was used to remove batch effects.

Principal component analysis (PCA) was performed with in R using the prcomp function based on top 500 highly variable genes. t-distributed stochastic neighbor embedding (t-SNE) was performed with the Rtsne (version 0.13) based on top 500 highly variable genes. The correlation between bulk RNA-seq and single cell samples was calculated by log_2_(TPM+1). All the figures were plotted by ggplot2 in R.

Pseudotime analyses were performed by the Monocle R package based on top 500 highly variable genes. The bulk RNA-seq correlation with pseudotime was calculated between RNA-seq log_2_(TPM+1) and simulated expression score of pseudotime order.

## Acknowledgments

We thank Ellen van den Bogaard and Matthijs Gerritse for discussion and for providing technical support, Eva Janssen-Megens, Siebe van Genesen and Rita Bylsma for operating the Illumina analyzer and initial data output. We thank the Studer Lab (Sloan Kettering Institute, New York) and the ENCODE Consortium for sharing their data. This research was supported by Netherlands Organisation for Scientific Research (NWO/ALW/MEERVOUD/836.12.010, HZ); Radboud University fellowship (HZ); Brazilian Science without Borders program (ES) and Chinese Scholarship Council grant 201406330059 (JQ).

## Author Contributions

ES, QX, QL, FT and HZ conceived and designed the experiments and wrote the manuscript. ES, QL, JQ, HHMR, KB, IP, WMRvdA and DA performed the experiments and provide reagents. ES, QX, YZ and HZ analyzed the data.

## Supplementary material and methods

### EXPERIMENTAL MODEL AND SUBJECT DETAILS

Human embryonic stem cells (ESC), human induced pluripotent stem cells (iPSC), and human primary keratinocytes (KCs) were cultured under the conditions described in method details.

### METHOD DETAILS

#### Human iPSCs generation and cell lines maintenance

Dermal fibroblasts of two health control individuals were used to generate induced pluripotent stems cells (iPSCs). The reprogramming was performed by lentiviral transduction of human *OCT4, SOX2, KLF4* and *cMYC* factors using the hOKSMco-idTomFRT plasmid (Warlich et al., 2011). The reprogrammed cells were initially cultured on a layer of Mouse Embryonic Fibroblasts (MEFs) in embryonic stem cell medium Dulbecco’s Modified Eagle’s Medium/Ham’s Nutrient Mixture F12 (DMEM/F12; GIBCO), 20% knock-out serum replacement (KSR; GIBCO), 2mM L-Alanyl-L-Glutamine (Sigma-Aldric), 100μM Non-Essential Amino Acids (NEAA; Sigma-Aldrich), 0.1 mM betamercaptoethanol (Sigma-Aldrich), 10ng/mL FGF2 (Sigma-Aldrich) and Penicillin Streptomycin (P/S, GIBCO). They were grown 37°C in 5% CO2. The cells were passaged every seven days by cutting the colonies in smaller pieces with a needle. After the cutting the cells were scraped off the surface with a pipette tip and transferred to a new MEF coated dish. iPSCs carrying *TP63* mutations were described previously (25).

All iPSC lines were later adapted into Essential 8™ medium (GIBCO) and more recently into Essential 8™ Flex Medium (GIBCO), following the guidelines from the manufacture. Basically, human iPSCs (niPSC1 and niPSC2, R204W, R304W and iso-R304W) and ESC-H9 (Thomson et al., 1998) (collectively called as PSCs) were cultured in dishes coated with 0.5 μL/cm^3^ Vitronectin (VTN-N) Recombinant Human Protein (GIBCO). The VTN-N was diluted 1:100 using Dulbecco’s Phosphate-Buffered Saline (DPBS, GIBCO). Cells were cultured in 2 mL/well (6-wells plate) with Essential 8™ Flex basal medium (GIBCO) with 2% supplement and 1% Penicillin Streptomycin (GIBCO) (E8 Flex++ medium). The medium was according to the manufactures’ instructions, and dishes were incubated at 37°C and 5% CO_2_.

Primary human keratinocytes (KCs) of a healthy control (control KCs) and patients carrying *TP63* mutations; R204W and R304W were described previously (67). Cells were cultured in 6 or 12 well plates in Keratinocyte Basal Medium (KGM, Lonza) supplemented with 100 units/mL P/S (GIBCO) 0.1 mM ethanolamine (Sigma-Aldrich), 0.1 mM O-phosphoethanolamine (Sigma Aldrich) 0.4% bovine pituitary extract, 0.5 μg/mL hydrocortisone, 5 μg/mL insulin and 10 ng/mL epidermal growth factor (all Lonza cat # 4131). All derived PSCs and KCs were checked for mycoplasma contamination periodically.

#### Differentiation of human PSCs into iKCs

PSCs were seeded as single cells before the initiation of differentiation. In details, PSCs were washed twice with 2 mL DPBS and detached by incubating 10 min with 1 mL/well (6-wells plate) StemPro™ Accutase™ Cell Dissociation Reagent (Accutase, GIBCO). Next, cells were added to 5 mL DMEM/F12 (GIBCO) and centrifuged for 5 min at 300 × *g*. Afterwards, supernatant was removed and cells were washed with 5 mL DMEM/F12. After centrifugation for 5 min at 300 × *g*, the pellet was re-suspended in 2 mL E8 Flex++ medium supplemented with 1× RevittaCell supplement (GIBCO). Cells were seeded in culture dished coated with Geltrex™ LDEV-Free Reduced Growth Factor which were incubated 1 hour at 37°C in 5% CO_2_ before seeding.

One day after seeding, differentiation was induced by incubating cells in keratinocyte induction medium (keratinocyte serum free medium (KSFM, GIBCO) + 0.2 ng/mL Epidermal Growth Factor (EGF) + 30 μg/mL Bovine Pituitary Extract (BPE) (both on GIBCO cat#17005-042) + 1% Penicillin Streptomycin (GIBCO) + 10 ng/mL Bone Morphogenetic Protein-4 (BMP-4, PromoKine) + 0.3 mg/mL retinoic acid (RA, Sigma-Aldrich). After 7 days cells were split and seeded in Geltrex coated plates with induction medium, and kept in induction medium until day 15. From day 15 to day 30 cells were cultured in KSFM (GIBCO) + 0.2 ng/mL EGF + 30 μg/mL BPE (both on GIBCO cat#17005-042) + 1% P/S (GIBCO). In the experiments for testing small molecule compounds, cells were cultured from day 7 in KSFM (GIBCO) + 0.2 ng/mL EGF + 30 μg/mL Bovine Pituitary BPE (both on GIBCO cat#17005-042) + 1% P/S (GIBCO) (with 0.3 mg/mL RA + 10 ng/mL BMPA-4, until day 15, and without RA and BMP-4 after day 15) supplement with either 1mg/mL Heparin Sodium Salt (Heparin, Sigma-Aldrich), 0.25uM Valproic Acid (VA, Sigma-Aldrich) or 100uM Suramin (Santa Cruz), all diluted in basal KSFM (GIBCO). The medium was refreshed every other day and dishes were incubated at 37°C and 5% CO_2_.

#### 2D differentiation of iKCs

The iKCs were induced in the basal keratinocyte medium KSFM (GIBCO) + 0.2 ng/mL EGF + 30 μg/mL Bovine Pituitary BPE + 1% P/S (GIBCO) with 10% of Fetal Bovine Serum (FBS) (GIBCO), when cells were at 60% confluency. Coverslips and RNA were collected prior induction, 0 hours as well 48 and 72 hours after induction.

#### Three germ layers differentiation

Human iPSC generated in this study were characterized for their differentiate potential into the three germ layers using Human Three Germ Layer Kit (R&D systems),according to the manufactures’ instructions with minor modifications. Briefly, single cell seeding was performed as described above, instead of clumps as described in the instruction of the kit (Figure S1). 24 hours after seeding, cells were induced with RPMI 1640 Medium, GlutaMAX (GIBCO) medium with differentiation supplements (for ectodermal, mesodermal or endodermal). RNA was collected after 2-3 days after the mesodermal induction, 3 days after the ectodermal induction and 4 days after the endodermal induction. Pluripotency characterization methods for R204W and R304W iPSCs were previously described (25).

#### Generation of lentiviral constructs, virus production and transduction

Lentiviral constructs were made through MultiSite Gateway 3-Fragment Recombination Reaction (Invitrogen). pEntry221-EFL-promoter and pEntry201-p63mutantR304W (kindly provided by D. Aberdam) were recombined into the 2K7bsd lentivector (68) using LR Clonase II Plus Enzyme Mix (Invitrogen). Lentiviruses carrying p63 mutant R304W were produced in HEK293T cells using Lipofectamine™ 3000 Transfection Reagent (Invitrogen, L3000015) according to the standard protocol. Virus was collected and concentrated about 100-fold by ultracentrifuge. For transduction, cells were cultured in E8++ medium and 50μL of p63 R304W mutant virus was added. Lentivirus-transduced cells were washed with 1x PBS 24 hours post-transduction and cultured in E8++ medium in presence of 6 μg/ml Blasticidin S HCl (GIBCO) for at least 2 weeks before starting differentiation.

#### RNA extraction and qRT-PCR

Total RNA was isolated at day 0 or 4, 7, 15 (WT and EEC-iPSC) and 30 (for WT only) days after the induction of epidermal commitment. Cell aggregates were lysed on plate using 350 μL RA1 (NucleoSpin^®^ RNA kit (Macherey-Nagel) buffer with 3,5 μL β-mercaptoethanol (Sigma-Aldrich) and snap-frozen in liquid nitrogen. Total RNA was extracted with NucleoSpin^®^ RNA kit (Macherey-Nagel) and converted into 100-500ng cDNA using the iScript cDNA synthesis kit (Bio-Rad). RT-PCR was performed with 10–100 ng of cDNA template in a 25 μl total reaction volume (12.5 μl iQ SYBR Green Supermix (2×) (Bio-Rad), 0.5 mM of each gene-specific primer, H_2_O up to volume). Duplicates of each sample were ran on a Bio-Rad CFX-96 real-time PCR system: denaturation at 95°C, 40 cycles of 15 seconds at 95°C, 45 seconds at 58°C, followed by a melting curve of 1 minute at 95°C, 1 minute at 65°C and 10 seconds at 65°C. Primers used for the reaction (Supplementary table 9) were designed with Primer3 http://bioinfo.ut.ee/primer3-0.4.0/). Glucuronidase Beta (*GusB*) was used as housekeeping gene to normalize cDNA levels.

#### Statistical analysis

The significance of the qRT-PCR results was determined by Student’s t-test and represented as mean ± SEM followed by a Bonferroni post-hoc test with the software Graphpad Prism 6.0. P-values below 0.05 were considered significant and are indicated in the figures.

#### Immunostaining

Cells were seeded in Geltrex (GIBCO) coated coverslips. At different days of differentiation coverslips were harvested. Firstly, they were washed in 1× PBS and fixed in 4% paraformaldehyde/1 × PBS for 15 min at room temperature or in methanol (Sigma-Aldrich) for 20min at −20C. After a 1 × PBS wash, cells were incubated in blocking buffer containing (0.01% Triton X-100, 5% normal goat serum (NGS) in 1× PBS) for 1 hour. Later cells were incubated with primary antibodies (Key resource table) diluted in antibody dilution buffer (10% bovine serum albumin (BSA, Sigma-Aldrich) overnight. Slides were mounted in VECTASHIELD Antifade Mounting Medium with DAPI (Vectorlabs). Images were acquired in a Leica DM Fluorescence Microscope.

#### Bulk RNA-sequencing

The NuGEN Ovation RNA-Seq (version 2, NuGEN) protocol was carried out on 100 ng of initial RNA. Ribosomal RNA (rRNA) was removed using targeted depletion with the NuGEN Ovation RNA-Seq platform (NuGEN) using the InDA-C Technology. In brief, total RNA is reverse transcribed using oligo-d(T) and random primers. cDNA fragments went through end repair, ligation, strand selection and adaptor cleavage following the manufacture’s recommendations. RNA-seq libraries were amplified using 10 cycles. The purified PCR products were then quantified using an Agilent bioanalyzer (DNA-1000 kit). We selected 300bp fragments using E-Gel™ SizeSelect™ II Agarose Gels, 2% (E-Gel, GIBCO). DNA fragments from the complete and size-selected library were detected and confirmed, respectively, using an Agilent bioanalyzer (DNA-1000 or High sensitivity kit). Ready to sequence RNA-seq libraries (size-selected) were tested for the expression of lineage-specific genes and quantified with a KAPA library quantification kit (KAPA Byosystems). All sample libraries were prepared were sequenced on Illumina HiSeq 2500 (Illumina, San Diego, CA, USA) for paired-end reads of 75×75bp.

#### Bulk RNA-sequencing analysis

RNA-seq reads were aligned to the reference genome (human hg19) using hisat2 (version 2.1.0) with default parameters. The resulting SAM files were sorted with read names and converted to a BAM file using SAMtools (version 1.7). The deepTools bamCoverage (version 3.0.2) tool was used to generate bigwig files with default parameters. Read counts representing gene expression levels were calculated with HTSeq (version 0.9.1) using default parameters and Genecode database annotations (version 19). Only protein coding genes were retained for the downstream analyses. Differential expression analyses were performed using DESeq2 (version 1.18.1) with adjusted p-value < 0.05 as the cutoff. The TPM (transcripts per million) values were calculated with the Scatter R package. Principal component analysis (PCA) was performed with the R programming language prcomp function based on top 500 highly variable genes. t-distributed stochastic neighbor embedding (t-SNE) was performed with the Rtsne (version 0.13) package in R based on top 500 highly variable genes. The top variable genes were decided by the variance of centered and scaled log_2_(TPM+1) among all samples. All Gene Ontology (GO) analyses were performed with ToppGene method (https://toppgene.cchmc.org/), and only Biological Process terms were kept for downstream analyses. The top four terms with p-value of each cluster were chosen. All the terms with log 10(p-value) < 4 and gene-ratio < 0.1 were then removed for the final plots. All the p-value are −log (raw.p.value,10). All PCA and tSNE scater plots and GO-term results were generated with the ggplot2 (version 2.2.1) package. All the heatmaps were generated using the ComplexHeatmap version 1.17.1) package with centered and scaled log_2_(TPM+1) score.

#### Single cell RNA sample preparation

A modified STRT-seq protocol was for the generation of the single-cell transcriptome profiles. The main steps are described below.

##### Single cell isolation

Cells were detached with Accutase in single-cell suspensions as described above. Cells were added to drops of 1% BSA on 1× DPBS (GIBCO) in a cell culture dish and transferred from one drop to another to wash the cells. Cells that looked phenotypically bright and round were then randomly picked with a mouth pipet.

##### Lysis and reverse transcription

Isolated cells were transferred to 200-μL PCR tubes containing 2.55 μL lysis buffer (0.05 μL RNase inhibitor (40 U μl^−1^, Takara), 0.095 μL 10% Triton X-100 (Sigma-Aldrich), 0.5 μL dNTP (10 mM), 0.1 μL including the external RNA controls consortium (1:2,000,000 dilution of ERCC RNA Spike-In Mix, Invitrogen), and 1.305 μL Nuclease-free water) and 0.5 μL oligo(dT) barcode primers (10uM, 5′-TCAGACGTGTGCTCTTCCGATCTXXXXXXXXNN NNNNNNT25-3′, where X8 represents predesigned 8bp cell-specific barcodes and N8 represents 8nt random unique molecular identifiers (UMIs)). Next, tubes were vortexed for 40s and incubated at 72°C for 3 min to release RNA molecules. Reverse transcription was performed by adding 2.85 μL of the reverse transcription mix (0.25 μL SuperScript II reverse transcriptase (200U μL-1, Invitrogen), 0.125 μL RNase inhibitor (40 U μl^−1^, Takara), 1 μL SuperScript II first-strand buffer (5×), 0.25 μL DTT (0.1 M) (both from Invitrogen cat# 18064022), 1 μL betaine (5 M), 0.03 μL magnesium chloride (1 M), 0.05 μL TSO (100 μM, 5′-AAGCAGTGGTATCAACGCAGAGTACATrGrG+G-3′; ‘rG’, riboguanosines; ‘+G’, locked nucleic acid (LNA)-modified guanosine) and 0.145 μL nuclease-free water per tube). The mixture was incubated in the BIORAD T100™ Thermal Cycler (25°C for 5 min; 42°C for 60 min; 50°C for 30 min; 72°C for 10 min and hold at 4°C).

##### PCR amplification

After reverse transcription, cDNA was amplified by adding 7.5 μL of the PCR mix (6.25 μL of 2X KAPA HiFi HotStart ReadyMix (KAPA Biosystems), 0.125 μL IS PCR primers (10 μM, 5′-AAGCAGTGGTATCAACGCAGAGT-3′), 0.625 μL 3′P2 primers (10 μM, 5′-GTGACTGGAGTTCAGACGTGTGCTCTTCCGATC-3′) and 0.5 μL nuclease-free water to each PCR tube. Samples were mixed and incubated in the Thermal Cycler (95°C for 3 min; 4 cycles of: 98°C for 20 s, 65°C for 30 s, 72° for 5 min; 14 cycles of: 98°C 20 s, 67°C for 15 s, 72°C for 5 min; then 72°C for 5 min; hold on 4°C).

##### Pooling and purification

25 cells with different cellular barcodes were combined into one sample. For this, 6.25 μL, half of the PCR products (13 μL) were pooled together and purified by using MiniElute^®^ spin columns (QIAGEN). For the PCR purification, 5x PB (on QIAGEN cat# 28104) buffer was mixed with the pooled samples, added to the spin column and centrifuged for 1 min at 13.3 × 10^4^ rounds per minute (rpm). 700 μL PE (on QIAGEN cat# 28104) buffer was then added per spin column, which were subsequently centrifuged two times 1 min at 13.3 × 10^4^ rpm. The cDNA was eluted by adding 51 μL nuclease-free water, incubated 1 min at 50°C and centrifuged 1 min at 13.3 × 10^4^ rpm.

After the column purification, samples were purified twice using Agencourt AMPure XP (Beckman Coulter). For the XP bead purification, 0.8× XP beads were added to each sample, mixed, and incubated for 15 min at room temperature (RT). After a short spin, samples were placed onto a magnetic rack for 5 min. The supernatant was discarded and beads were washed with 200 μL 80% ethanol (Sigma-Aldrich) twice (ethanol was added and incubated for 30 s before discarding). After the second wash, all ethanol was removed and the beads were dried for 5 min on air. DNA was finally eluted in32.5 μL nuclease-free water by incubated 3 min at RT. The tubes were placed back into the magnetic rack and the clear supernatant (containing cDNA) was transferred to a new tube.

##### Biotin primer PCR

In this protocol, biotin was used to enrich for the 3′ end cDNA fragments. Attachment of biotin to these fragments was performed by PCR with biotin-linked primers (linkage at 3′ end). For the attachment with the biotin, 30 ng cDNA and the PCR mix (25 μL of 2× KAPA HiFi HotStart ReadyMix (KAPA Byosystems), 2 μL IS PCR primers (10 μM), 2 μL of one of the seven biotin index primers (10μM) and 21 μL nuclease-free water– 30 ng cDNA) were mixed and incubated in the Thermal Cycler (95°C for 3 min; 5 cycles of: 98°C for 20 s, 67°C for 15 s, 72°C for 5 min; then 72°C for 5 min; hold at 4°C). The PCR products were purified twice with 0.8× XP beads as previously described.

##### DNA fragmentation

About 400 ng of DNA were sheared to approximately 300bp fragments with Diagenode Bioruptor^®^ at 4°C (PICO ON: 30 s; OFF: 30 s; 12 cycles). After sonication, purification with XP beads was performed as described before. After purification, DNA was eluted in 50 μL elution buffer (EB, on QIAGEN cat# 28104).

##### Biotin enrichment

DNA fragments attached to biotin were enriched by using Dynabeads™ MyOne™ Streptavidin C1 beads (Invitrogen). For this enrichment, C1 beads were vortexed and 10 μL per sample were added to a new PCR tube. Next, 200 μL of 2× binding and washing (B&W) buffer (10 mM Tris-HCl, pH 7.5, 1 mM EDTA, 2.0 M NaCl) were added to the C1 beads, vortexed, and placed in a magnetic rack for 5 min. After incubation, the supernatant was discarded and the beads were eluted in 50 μL EB. The samples were mixed with the beads and were incubated in a rotator (30 rpm) for 1 hour at RT. Next, the samples were placed onto a magnetic rack and supernatant was discarded. Beads were washed with 100 μL of B&W buffer, whereby the supernatant was removed after 5 min incubation on the magnetic rack at RT. The beads (with DNA attached to it) were eluted in 25 μL nuclease-free water.

##### Library preparation

We started library preparation with end repair and A-tailing. For this, 25 μL sample were mixed with 3.5 μL End Repair & A-Tailing buffer and 1.5 μL End Repair & A-Tailing Enzyme mix and incubated in the Thermal Cycler (20°C for 30 min; 65°C for 30 min; hold on 4°C). Immediately after end repair and A-tailing, reagents for adaptor ligation were added (to 30 μL of product: 2.5 μL MQ, 15 μL ligation buffer, 5 μL DNA ligase and 2.5 μL NEXTflex^®^ DNA Barcodes (Bioo Scientific) and the mixture was incubated in the Thermal Cycler (20°C for 30 min; hold at 4°C). To remove the ligation reagents, samples were put onto the magnetic rack for several minutes before the supernatant was discarded. The C1 beads were washed once with 200 μL EB and were eluted in 21 μL MQ. Amplification of the library was achieved by adding 25 μL of 2× KAPA HiFi HotStart ReadyMix (KAPA Byosystems), 2 μL QP2 primer (10 μM) and 2 μL NEB short universal primer (10 μM) to the 21 μL adapter-ligated library and incubated in the Thermal Cycler (98°C for 45 s; 8 to 10 cycles of: 98°C 15 s, 65°C for 30 s, 72°C for 30 s; then 72°C for 1 min; hold on 4°C). Library preparation was completed by placing the tubes onto the magnetic rack and transferring the supernatant into a new PCR tube. The collected supernatant was purified using 0.9× XP beads and were finally eluted in 30 μL MQ.

##### Concentration and fragment size

During single cell sample preparation, concentrations of the samples checked by the Qubit^®^ fluorometer according to the Qubit dsDNA HS Assay Kits user guide (Invitrogen). Fragment size was measured by Agilent 2100 Bioanalyzer before sonication, after sonication, and after library preparation. Fragment size measurement was performed as described in the Agilent High Sensitivity DNA Kit Quick Start Guide (Agilent).

##### Sequencing

Single cell RNA-seq libraries were sequenced by Illumina NextSeq™ 500 with a sequencing depth of 50 million reads for paired-end reads of 20bp for Read 1 and 64bp for Read 2. A total of 1,250 single cells) were sequenced.

#### Single cell RNA-seq analysis

##### Pre-analysis of scRNA-seq Data

All the single cell RNA-seq samples were sequenced by the paired-end method. The Read 2 of the paired-end reads was comprised of an 8-bp cell barcode followed by an 8bp UMI sequence before the ploy(A) tail sequence, and the corresponding Read 1 contained exon reads of genes. The Read 1 file was split by the corresponding cell barcode in Read 2. Low quality reads and ploy(A) tail were removed, and only reads more than 37bp as clean reads were kept. The clean reads of each single cell sample were then aligned to the reference genome (hg19 and ERCC for human) using hisat2 (version 2.1.0) with default parameters. The duplicated reads from the same transcript were removed based on the UMI information in corresponding Read 2. For each single cell sample, read counts representing gene expression levels were calculated using HTSeq (version 0.9.1) with default parameters. hg19 GTF annotation file was download from Gencode (version 19), and then merged with the ERCC gene annotation for read counting. Only protein coding genes and ERCC gene counts information was kept and continued for downstream analyses.

##### scRNA-seq data normalization

**E**xpression levels in all samples were normalized with the Scatter (version 1.5.0) R/Bioconductor package by log_2_(TPM+1). For all 1,250 sequenced single cells, the low quality samples were filtered with five metrics: mapping rate, library size, gene number, percentage of counts mapping to mitochondrial genes and to ERCC transcripts. The mapping rate threshold was defined by mapping rate above 30%. The library size and gene number threshold were defined by the median absolute deviation (MAD) of above 3, the percentage of counts mapping to mitochondrial genes and to ERCC transcripts was defined by the MAD of below 3. In total, 964 single cell samples were kept for downstream analyses. Based on the SC3 cluster package, the top 500 highly variable genes were used, and the RUVSeq R package was used to remove batch effect.

##### scRNA-seq Data Dimensional Reduction

Principal component analysis (PCA) was performed with the R programming language prcomp function based on top 500 highly variable genes identified by SC3. t-distributed stochastic neighbor embedding (t-SNE) was performed with the Rtsne (version 0.13) package in R based on top 500 highly variable genes identified by SC3. The correlation between bulk RNA-seq and single cell samples was calculated by log_2_(TPM+1). The marker genes expression in PCA plot were log_2_(TPM+1). All the figures were plot by ggplot2 in R programming language.

##### Pseudotime analyses

Pseudotime analyses were performed by the Monocle R package based on top 500 highly variable genes identified by SC3. The heatmap of simulate gene expression was simulated by pseudotime order. k-means method was used to cluster the simulated expression to 6 clusters. The bulk RNA-seq correlation with pseudotime was calculated between RNA-seq log_2_(TPM+1) and simulated expression score of pseudotime order. GO analyses were performed with ToppGene method (https://toppgene.cchmc.org/). The top five terms by p-value for each cluster were chosen. All the terms with Log10(p-value) < 4 and gene-ratio < 0.1 were then removed for the final plots. All the p-value are −log (raw.p.value,10).

## Supplementary Tables

**Supplementary table 1.xlsx All bulk RNA-seq samples expression TPM table.**

**Supplementary table 2.xlsx Normal bulk RNA-seq samples cluster and go terms.**

**Supplementary table 3.xlsx Normal cell RNA-seq samples PCA PC loadings.**

**Supplementary table 4.xlsx RNA-seq samples day30 to KC regulate genes and go terms.**

**Supplementary table 5.xlsx All bulk RNA-seq samples cluster and go terms.**

**Supplementary table 6.xlsx EEC mutations overlapping up/down-regulated genes and go terms.**

**Supplementary table 7.xlsx All single cell RNA-seq samples PCA PC loadings.**

**Supplementary table 8.xlsx Single cell RNA-seq samples pseudotime branch simulation cluster and go terms.**

**Supplementary table 9.**
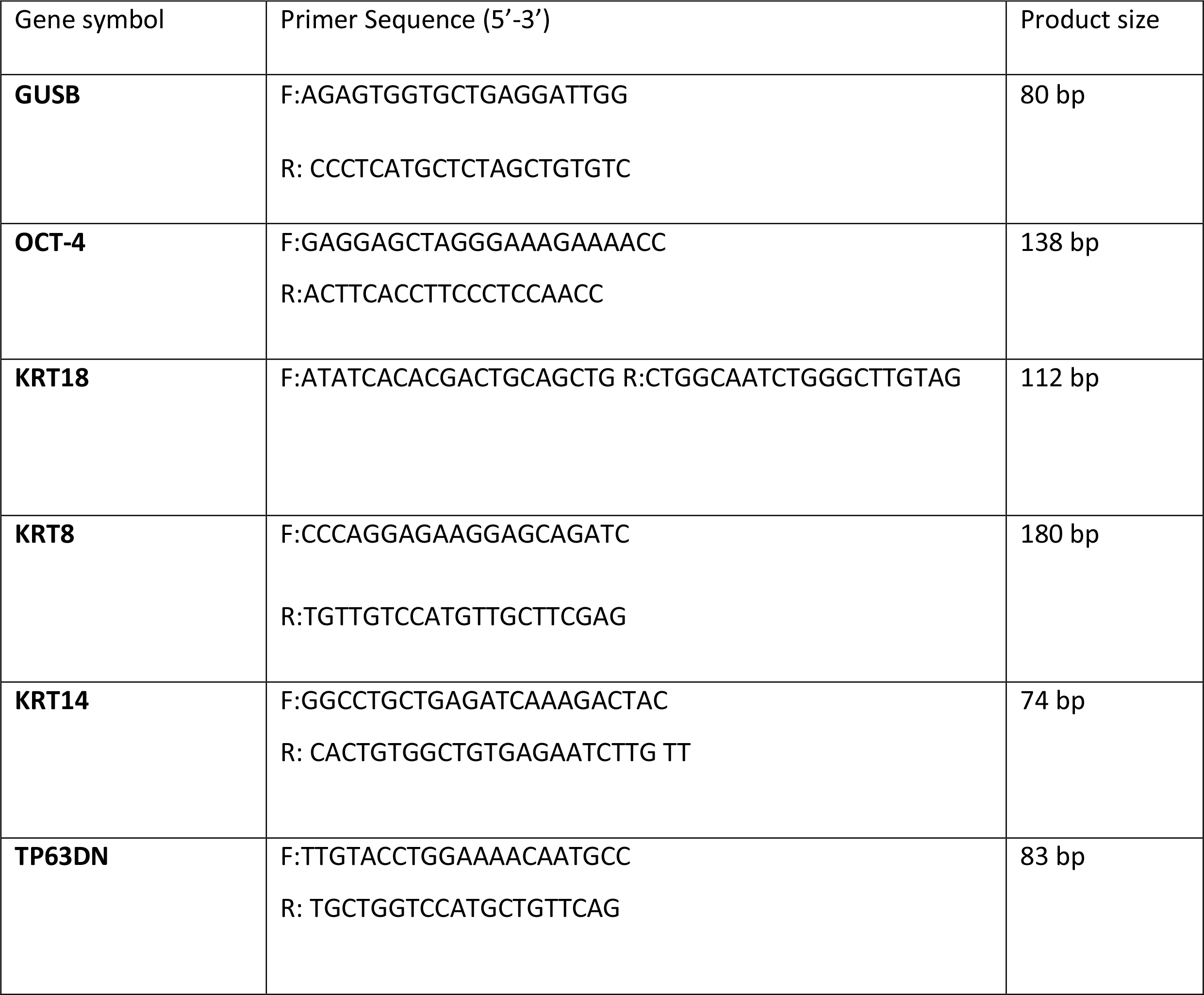

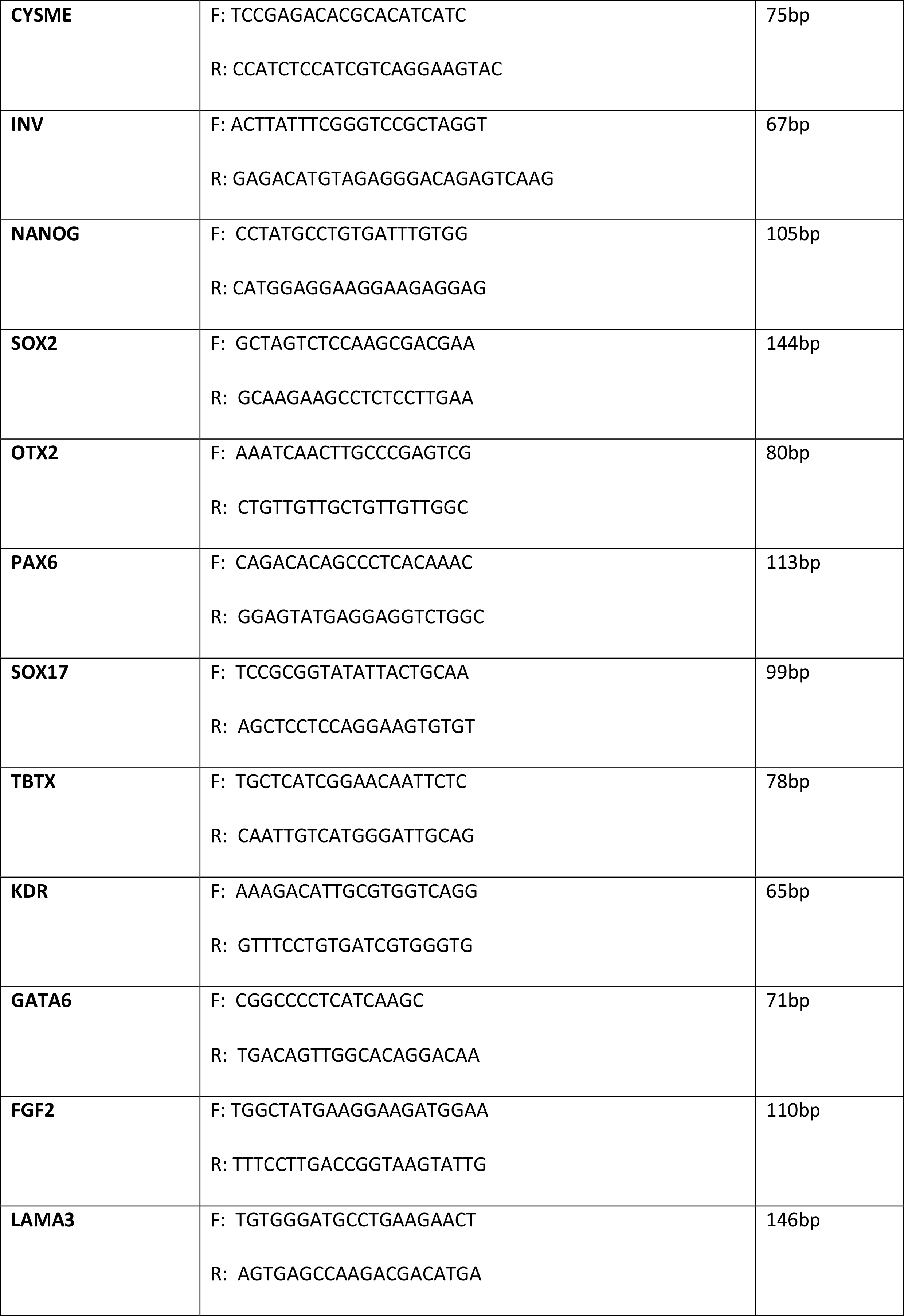

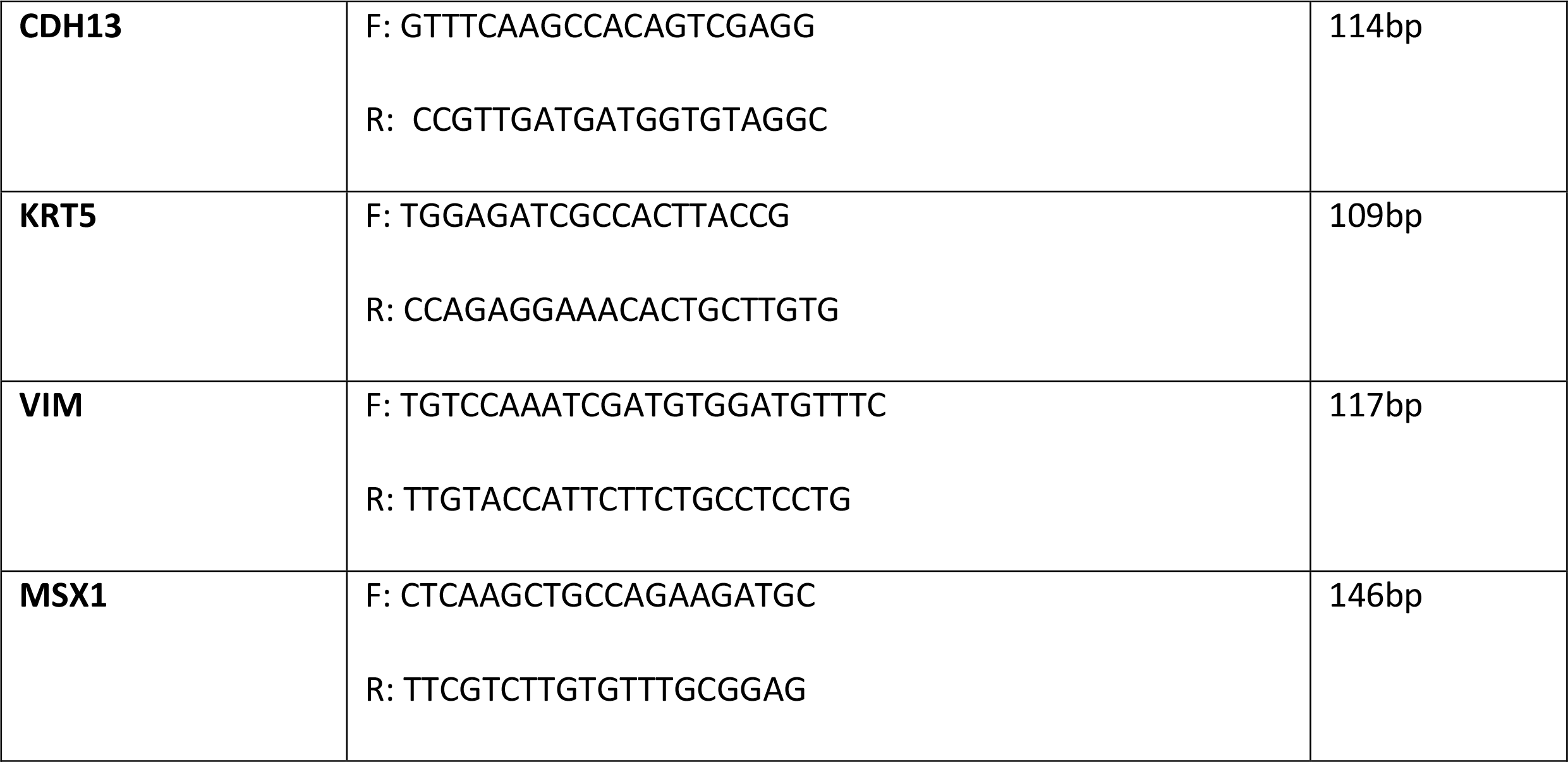

## Supplementary figure legends

**Figure S1.**
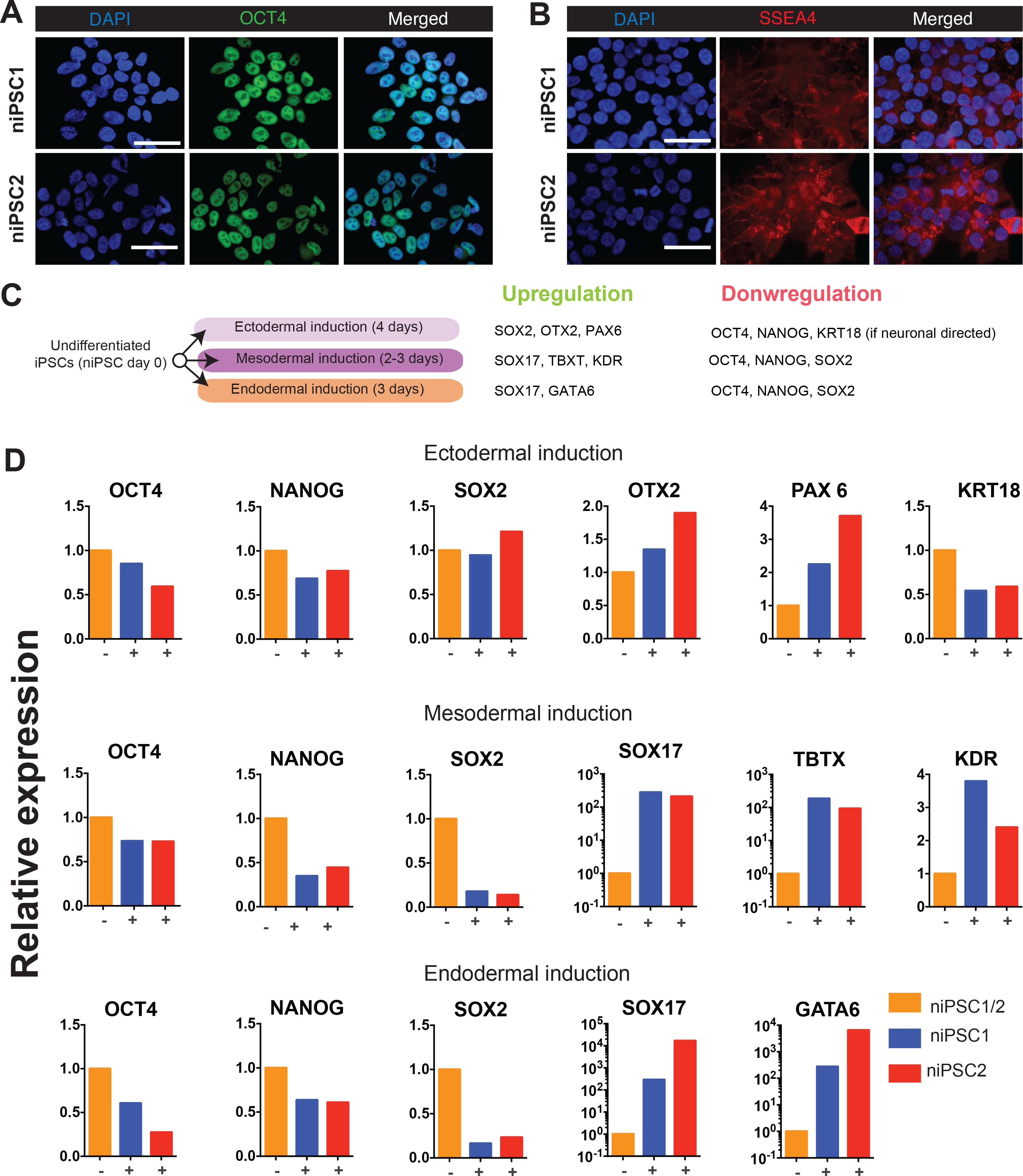
Pluripotency tests of control iPSCs. (A)Immunofluorescence staining of the pluripotency marker OCT4 (red) and DAPI (blue) in the newly generated control niPSC1 and niPSC2 at undifferentiated stage, day 0 (Scale bars, 50 μm). (B)Immunofluorescence staining for the pluripotency marker SSEA-4 (red) and DAPI (blue) in control niPSC1 and niPSC2 at undifferentiated stage, day 0 (Scale bars, 50 μm). (C)Scheme of the differentiation assay for differentiation potential towards three germ layers. Supplements for ectodermal, mesodermal or endodermal induction were added to basal medium. RNA was harvested at the indicated time points. Patterns of expected gene expression for each lineage specification is indicated on the right side. (D)qRT-PCR analysis of lineage-specific genes to confirm the differentiation potential of niPSC1 and 2 towards three germ layers. Gene expression is expressed as 2^- ΔΔCt values over GUSB and samples without supplement.

**Figure S2.**
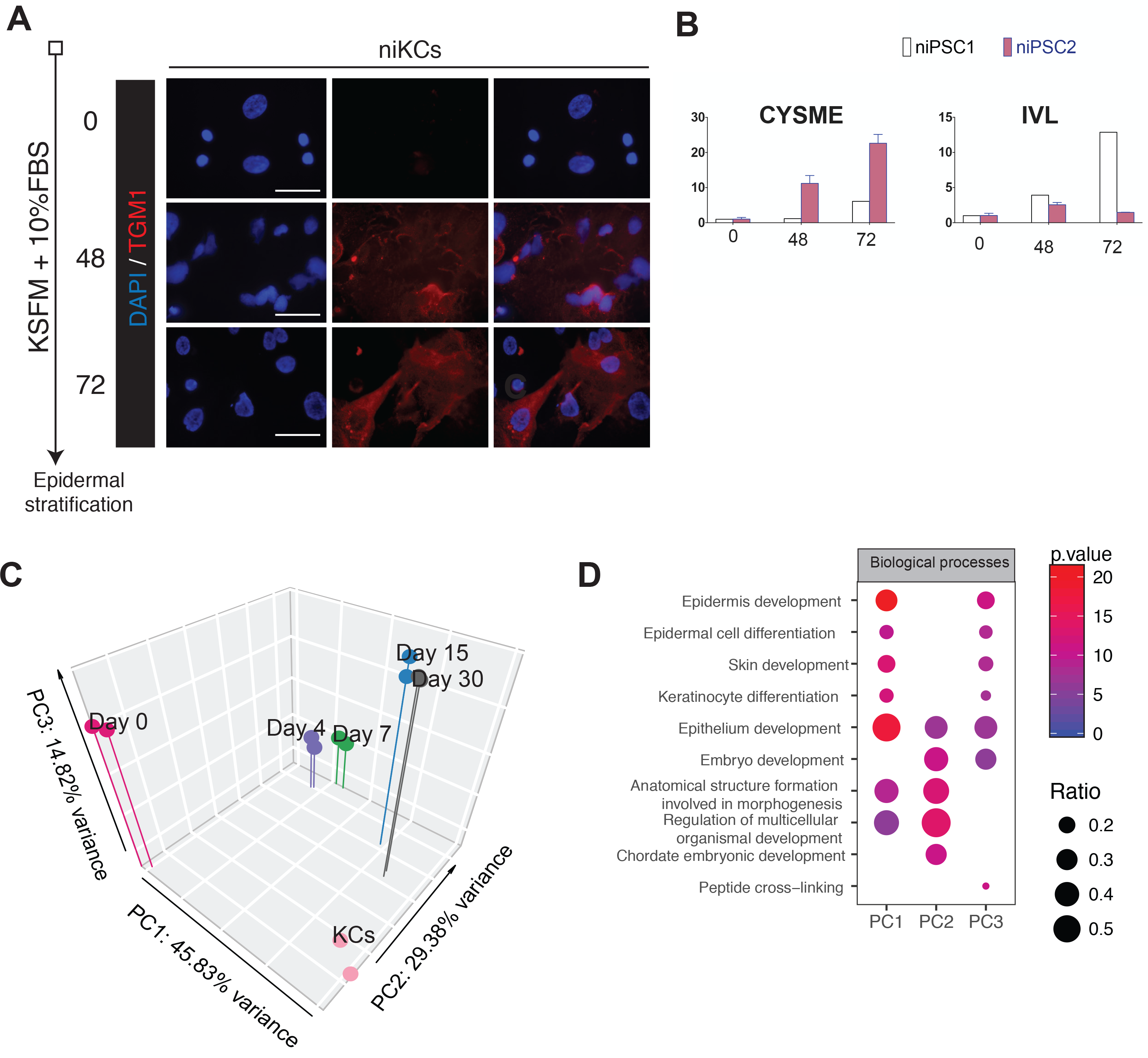
Characterization control PSC differentiation. (A)Terminal differentiation of day-30 iKCs into stratified epithelium was induced in a 2D culture system, with KSFM supplemented with 10% FBS, shown on the left. Upon induction, cells were analysed for the presence of the suprabasal marker transglutaminase 1 (TGM1) by immunostaining at 0, 48 and 72 hours (Scale bars, 50 μm). (B)qRT-PCR analysis of suprabasal epithelial markers CYSME and IVL. Samples were harvested at 0, 48 and 72 hours after induction. Gene expression is expressed as 2^- ΔΔCt values over GUSB and 0 hours samples (n=1 for niPSC1; n=2 for niPSC2). (C)3D PCA of transcriptomes of niPSC1 during epidermal differentiation and human primary keratinocytes. Different colors represent different days of differentiation. (D)Gene Ontology (GO) analysis showing biological process terms enriched for PC1, PC2 and PC3 axes. The gene ratio is indicated by the dot sizes and the significance by the color of the dot (Red: low pvalue: blue: high pvalue).

**Figure S3.**
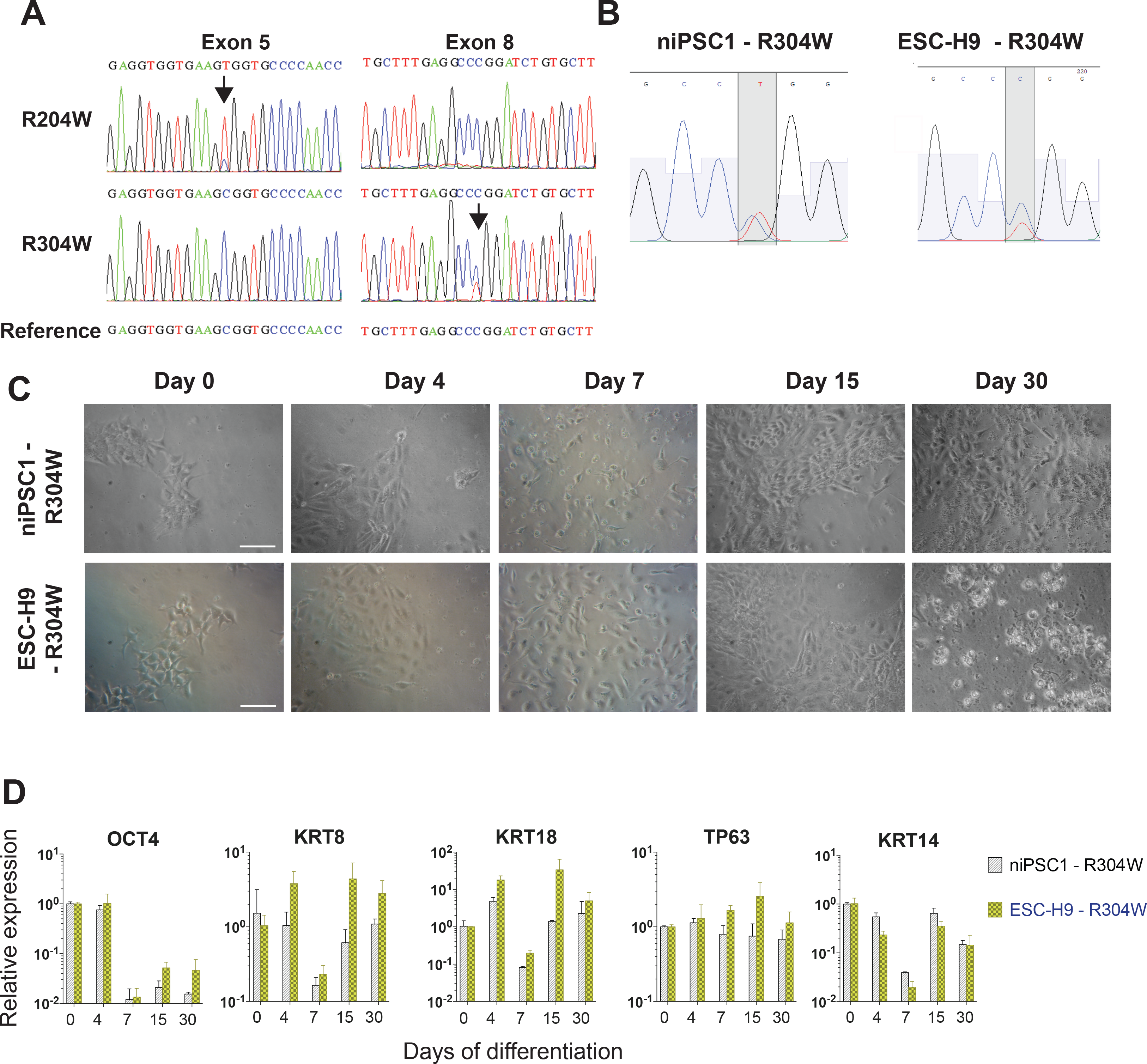
Validation and characterization of EEC-PSCs. (A)Sanger sequencing confirming the two heterozygous point mutations (a C>T substitution) in exon 5 (R204W) and exon 8 (R304W) in the *TP63* gene. Arrows indicate that the point mutations were detected on the specific exons, and reference sequences are shown at the bottom. (B)Sanger sequencing analysis of the presence of a C>T substitution in the mouse p63 sequence using mouse--specific primers. Two biological replicas in each cell line are shown. (C)Bright field images of two different isoR304W cell lines (niPSC1-R304W and ESC-H9 – R304W) throughout different days of differentiation (0,4,7, 15 and 30) (Scale bars, 100 μm). (D)qRT-PCR analysis of the pluripotency marker *OCT4*, simple epithelium markers *KRT8*, *KRT18* and epidermal markers *TP63* and *KRT14* in the two isoR304W cell lines. Gene expression is expressed as 2^- ΔΔCt values over *GUSB* and undifferentiated day 0 samples (n=2 per cell line).

**Figure S4.**
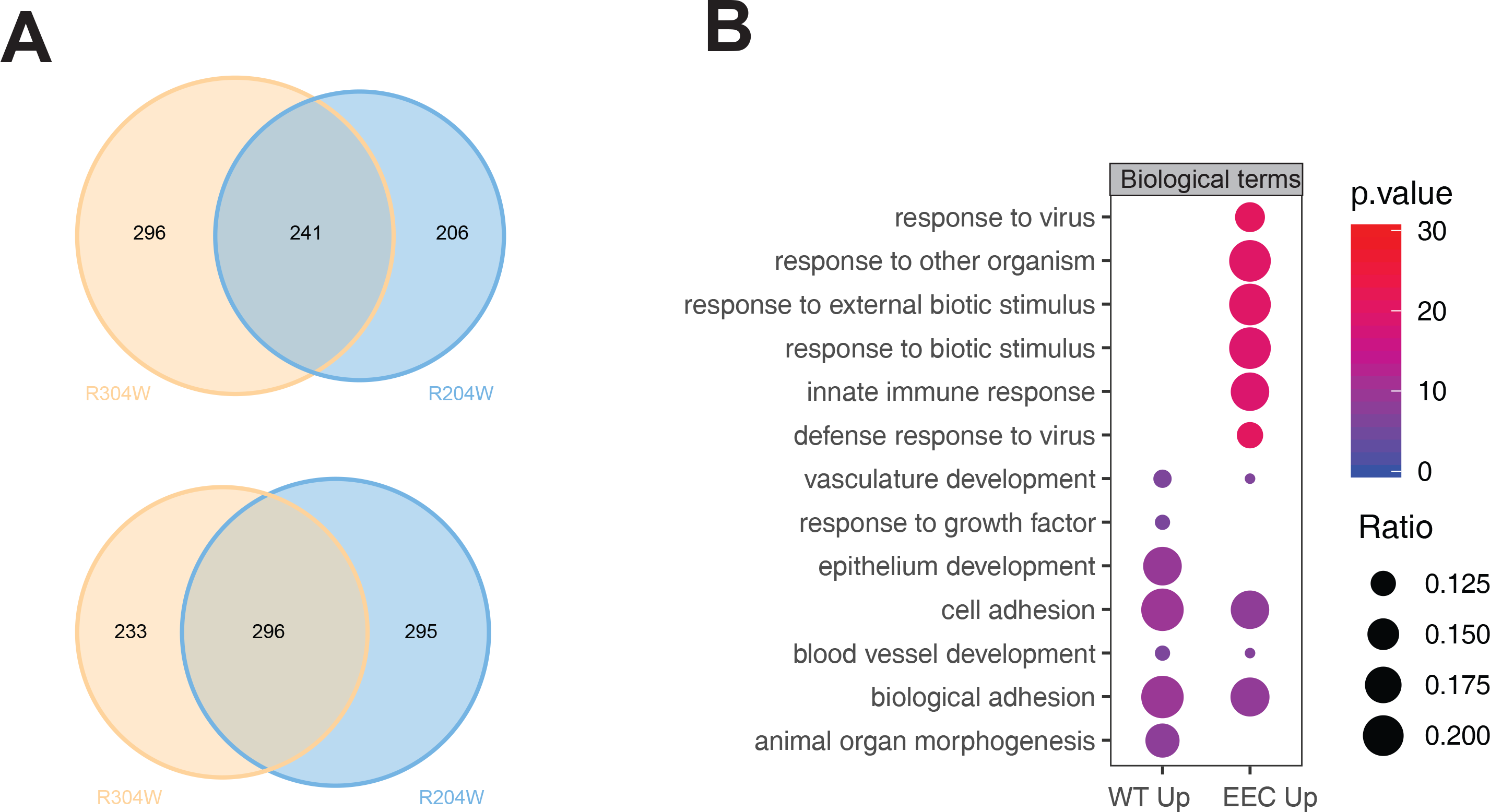
Consistently deregulated genes of EEC-PSCs on differentiation day 15. (A)Venn diagram showing the number of overlap genes from the pairwise comparison between control and EEC-iPSCs (R204W or R304W). The upper panel, downregulated in EEC-PSCs; the lower panel, upregulated in EEC-PSCs. (B)GO analysis showing biological process terms enriched for consistently regulated in both EEC-iPSCs. The gene ratio is indicated by the dot size and the significance by the color of the dot (Red: low pvalue: blue: high pvalue).

**Figure S5.**
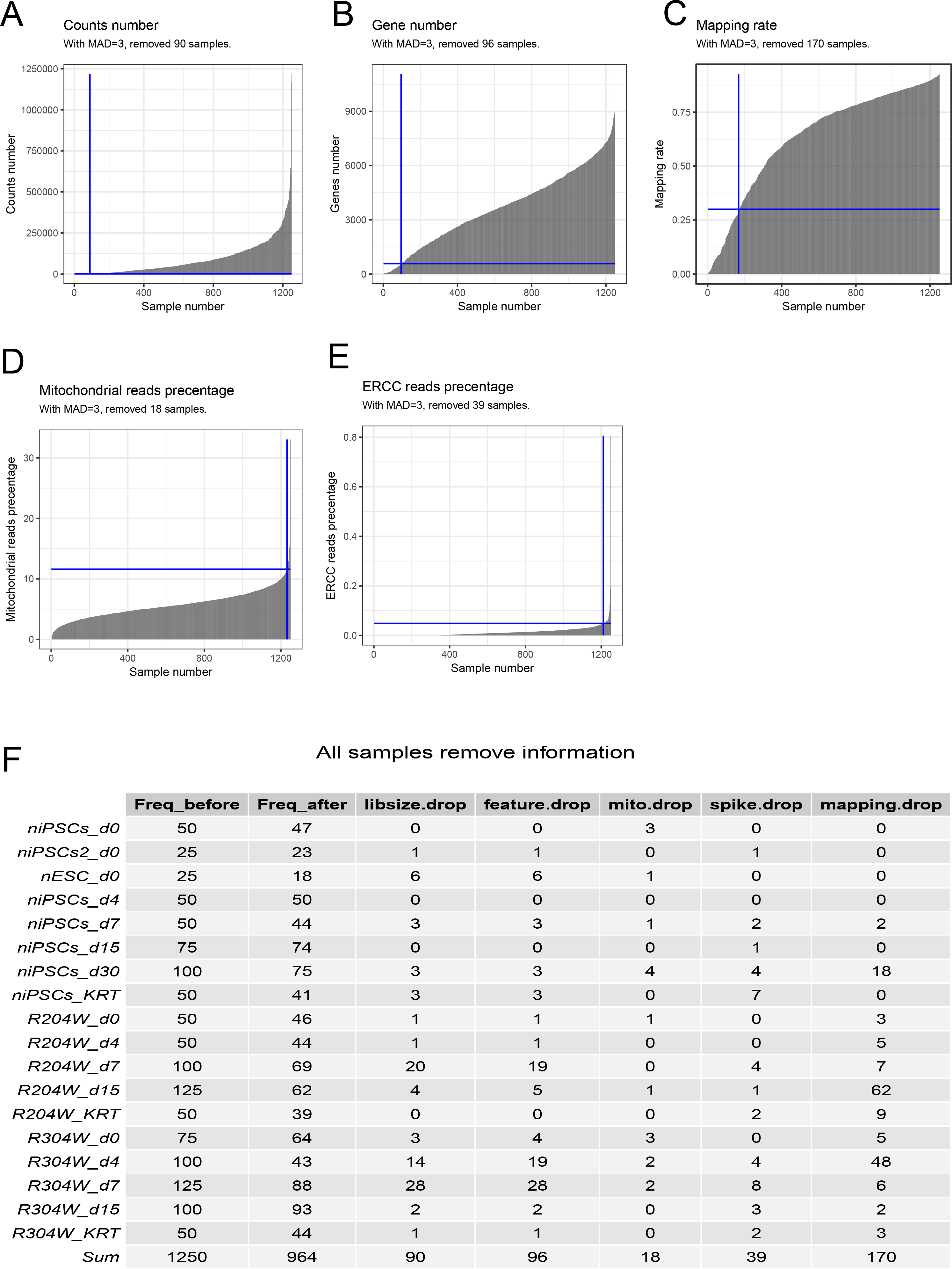
Quality checks of single cell RNA-seq. (A)Single cell RNA-seq quality check of count number of each single cell sample. Using the threshold defined by the median absolute deviation (MAD) of above 3, 90 single cell samples was removed. (B)Using the threshold defined by MAD above 3, 96 single cell samples was removed. (C)Using the threshold defined by mapping rate above 30%, 170 single cell samples was removed. (D)Using the threshold defined by MAD below 3, 18 single cell samples was removed. (E)Using the threshold defined by MAD below 3, 39 single cell samples was removed. (F) The summary of all single cell RNA-seq samples in quality checks. The total number of cells is 1250, and 964 single cell samples remained after quality checkfor the downstream analysis.

**Figure S6.**
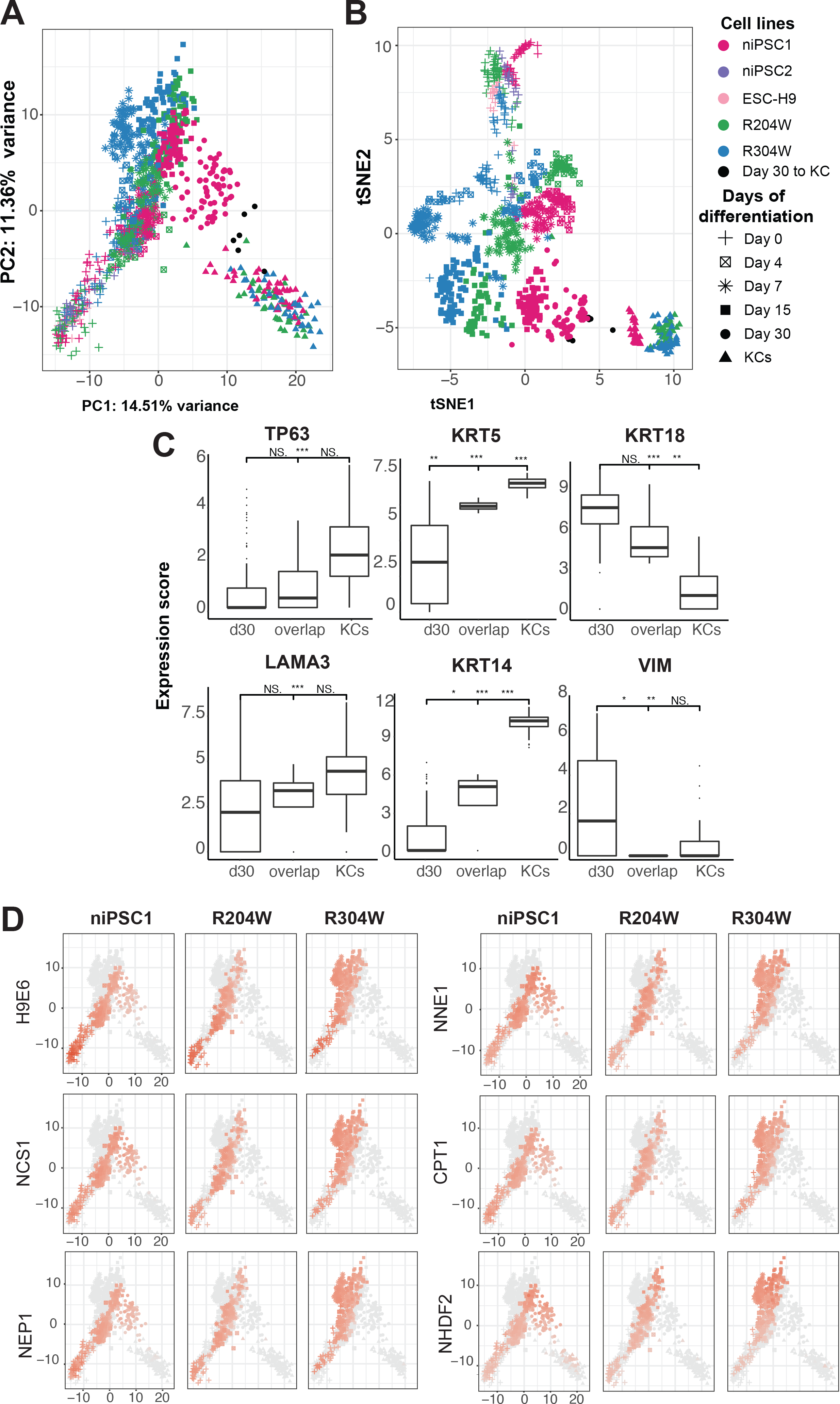
Heterogeneous populations of differentiating PSCs. (A)PCA and t-SNE analyses of the top 500 highly variable (HV) genes obtained from single-cell RNA-seq during epidermal differentiation of PSCs and primary keratinocytes. Colors represent cell lines; shapes represent differentiation days. The six cells that showed similar transcriptome to primary KCs are indicated in black. (B)Expression level (Log(TPM+1)) comparison of marker genes in the bulk day-30 iKCS, the six cells that showed similarity to primary KCs (black dots) and primary KCs. (C)Correlation of single-cell transcriptome against bulk RNA-seq data of cell types from diverse lineages shown in PCA (A). H9E6=human embryonic stem cells, NCS1= neural crest, NEP1= neural epithelial, NNE1=Non-neural Ectoderm, ESneu= Neuroectoderm and CPT1= Cranial Placode and non-embryonic: nhdf2=normal human dermal fibroblasts and PHNeu=?? cell types. Correlation coefficient is shown on the bar bellow the plots. Red: high correlation; gray: low correlation and other cell types.

**Figure S7.**
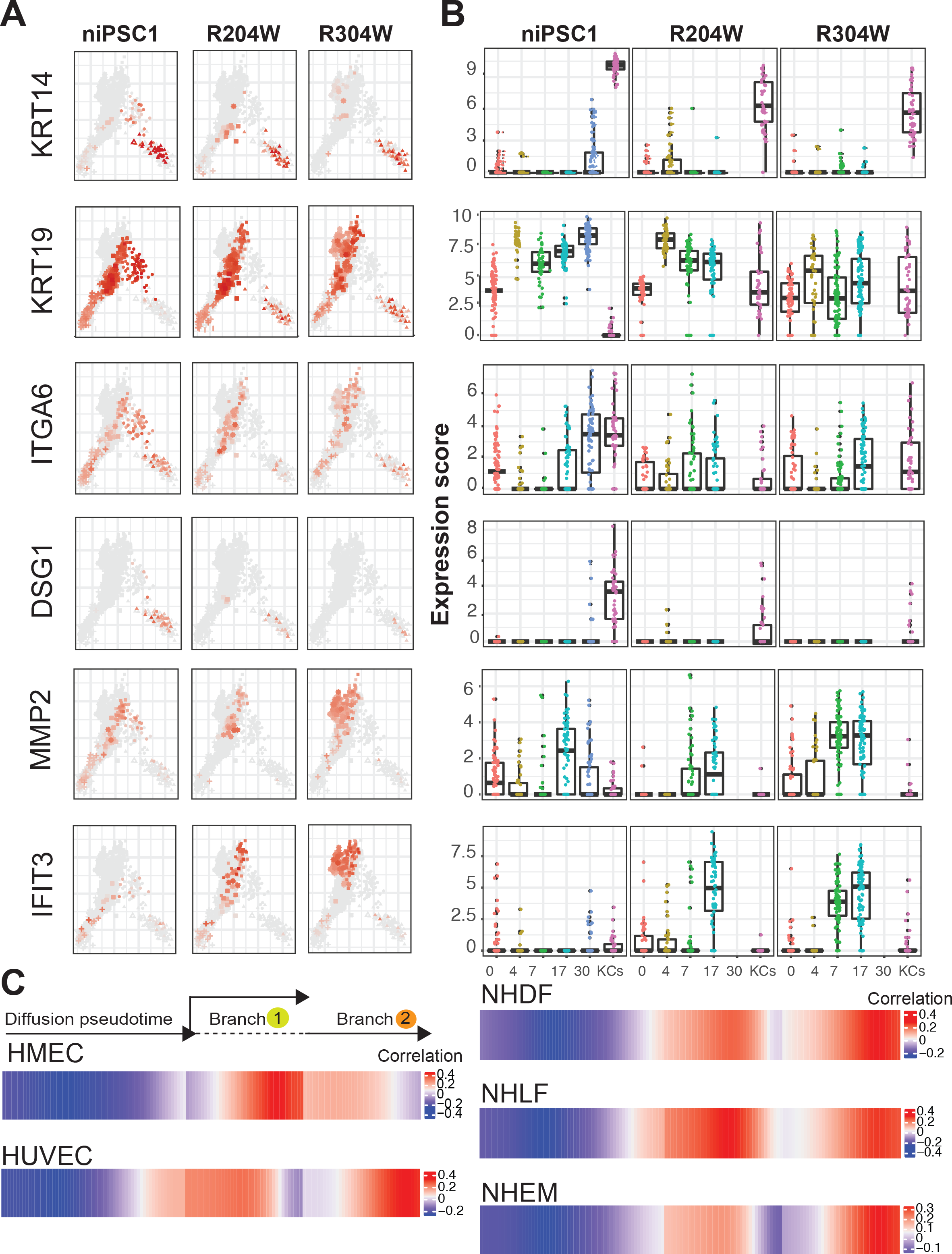
Enhanced mesodermal gene expression of differentiation PSCs. (A)Marker gene expression levels (Log(TPM+1)) in single-cell transcriptome shown in PCA. Epidermal/epithelial, *KRT14, KRT19, ITGA6* and *DSG1.* Non-epithelial, *MMP2* and *IFIT3.* Red: high correlation; gray: low correlation and other cell types. (B)Marker gene expression levels (Log(TPM+1)) in single-cell transcriptome shown by bar plots. Marker genes are labeled in panel (A).

